# High-throughput, microscopy-based screening, and quantification of genetic elements

**DOI:** 10.1101/2023.09.06.556327

**Authors:** Rongrong Zhang, Yajia Huang, Mei Li, Lei Wang, Bing Li, Aiguo Xia, Ye Li, Shuai Yang, Fan Jin

## Abstract

Synthetic biology relies on the screening and quantification of genetic components to assemble sophisticated gene circuits with specific functions. Microscopy is powerful tool for characterizing complex cellular phenotypes with increasing spatial and temporal resolution to library screening of genetic elements. Microscopy-based assays are powerful tools for characterizing cellular phenotypes with spatial and temporal resolution, and can be applied to large-scale samples for library screening of genetic elements. However, strategies for high-throughput microscopy experiments remain limited. Here, we present a high-throughput, microscopy-based platform that can simultaneously complete the preparation of an 8×12-well agarose pads plate, allowing for the screening of 96 independent strains or experimental conditions in a single experiment. Using this platform, we screened a library of natural intrinsic promoters from *Pseudomonas aeruginosa* and identified a small subset of robust promoters that drives stable levels of gene expression under varying growth conditions. Additionally, the platform allowed for single-cell measurement of genetic elements over time, enabling the identification of complex and dynamic phenotypes to map genotype in high-throughput. We expected that the platform could be employed to accelerate the identification and characterization of genetic elements in various biological systems, as well as to understand the relationship between cellular phenotypes and internal states, including genotypes and gene expression programs.

**Impact statement:** The high-throughput microscopy-based platform, presented in this study, enables efficient screening of 96 independent strains or experimental conditions in a single experiment, facilitating the rapid identification of genetic elements with desirable features, thereby advancing synthetic biology. The robust promoters identified through this platform, which provide predictable and consistent control over gene expression under varying growth conditions, can be utilized as reliable tools to regulate gene expression in various biological applications, including synthetic biology, metabolic engineering, and gene therapy, where consistent system performance is required.

## INTRODUCTION

In recent decades, genetic screens for visual phenotypes have been highly successful genetic approaches^1–3^. However, among the various platforms for mapping genotype to phenotype, microscopy-based methods are currently the only approach that allows for monitoring gene expression and/or cell behavior in single cells over time. This in-depth analysis has proven to be a potent tool for understanding gene circuit dynamics, cell behavior heterogeneity, and broad-spectrum phenotypes.^3–6^. It has become indispensable in biological engineering and synthetic biology, for applications such as library screening to develop new protein biosensors, synthetic enzymes, genetic elements, and circuits, signaling pathways, and multicellular communities^2, 7^. We have seen the development of microscopy-based, high-content assays, which allow for parallel monitoring of cells possessing different genotypes. The utilization of fluorescently labeled proteins or fluorescent markers can visualize cellular and subcellular phenotypes. This method can image millions of cells from thousands of genetic variants and visually assess their phenotypes in a pooled format^8–11^.

Recent advances in in situ sequencing methods allow for the visual reading of nucleic acid barcodes to map their genotypes^12–18^. When paired with Fluorescence-Activated Cell Sorting (FACS) ^19–22^, cells exhibiting desirable phenotypes can be isolated from pooled variants. However, these methods involve complex procedures, require expensive reagents, and need extensive computational pipelines. Using photoactivatable proteins to mark and sort cells necessitates additional genetic manipulation and high-resolution light patterning^19, 20^.

Microscopy-based approaches excel in analyzing the heterogeneous behavior of individual cells^6^, especially when screening for genetic elements with temporal or spatial properties, such as biosensor kinetics and gene expression dynamics^23^. Nonetheless, the preparation of samples for large-scale experiments has been a significant bottleneck in classical microscopy investigations, limiting the method’s throughput due to the labor-intensive nature of sample preparation. Even with the optimization of commercial multi-well dishes or cell microarrays for high throughput, the necessity of manual intervention to scan samples restricts the time resolution or the number of cells that can be analyzed^2^. Consequently, there is a pressing need for strategies capable of preparing hundreds of microscope samples simultaneously.

In our current study, we endeavored to augment the throughput of microscopy-based experiments by devising a sampling device that facilitates large-scale sample preparation. This tool is followed by a rapid, automated acquisition of images for all samples under the microscope. To demonstrate the versatility of our platform, we conducted a proof-of-concept study by screening and quantifying the activity of a library of natural promoters in *Pseudomonas aeruginosa*. Using the platform, we were able to simultaneously screen 96 promoter reporters in a high-throughput manner. We identified robust promoters that showed stable functions across a range of growth and stress conditions in *P. aeruginosa* and *Pseudomonas putida* KT2440. We further validated the robustness of the screened promoters by quantifying their temporal properties, such as growth rate, at the single-cell level. The 96-well agarose plate can be prepared using a standard protocol and is universally compatible with standard microscope object stages, making our platform easy to use for other groups. Because of the versatility and generality of our platform, we anticipate its wide utilization for accelerating the identification and characterization of genetic elements in a wide range of biological systems.

## RESULTS

### High-throughput microscopy-based platform for simultaneous analysis of hundreds of samples

We designed a high-throughput micro-sampling device for bacteria that can simultaneously complete the preparation of an 8×12-well agarose pads plate, thus facilitating the screening of 96 independent strains or experimental conditions in a single experiment (Figure 1, Step1 and Step3). The device employs a sandwich structure where bacterial cells are positioned between an agarose pad and a piece of cover glass. The thickness of the agarose pad is optimized to allow adequate air exchange while providing sufficient medium to foster bacteria growth (Figure S1, Methods section for detailed sampling procedures). Moreover, the 96-well agarose pad plate is specifically designed to be universally compatible with standard microscope object stages (Figure S2). When paired with a microscope equipped with a high-speed camera and Piezoelectric positioning stages, boasting response times of millisecond and microsecond ranges, our platform facilitates fast-scanning imaging of all 96 samples automatically within a mere 15 minutes (Figure 1, Step4). And with our streamlined experimental workflow, from sample preparation to one-shot image scanning (Figure 1, Step2 to Step4), the entire procedure can be completed within just 30 minutes, substantially boosting the screening throughput and efficiency. Finally, the resulting images were analyzed using custom-designed software programs or codes^24^ (Figure 1, Step5).

**Figure 1.**
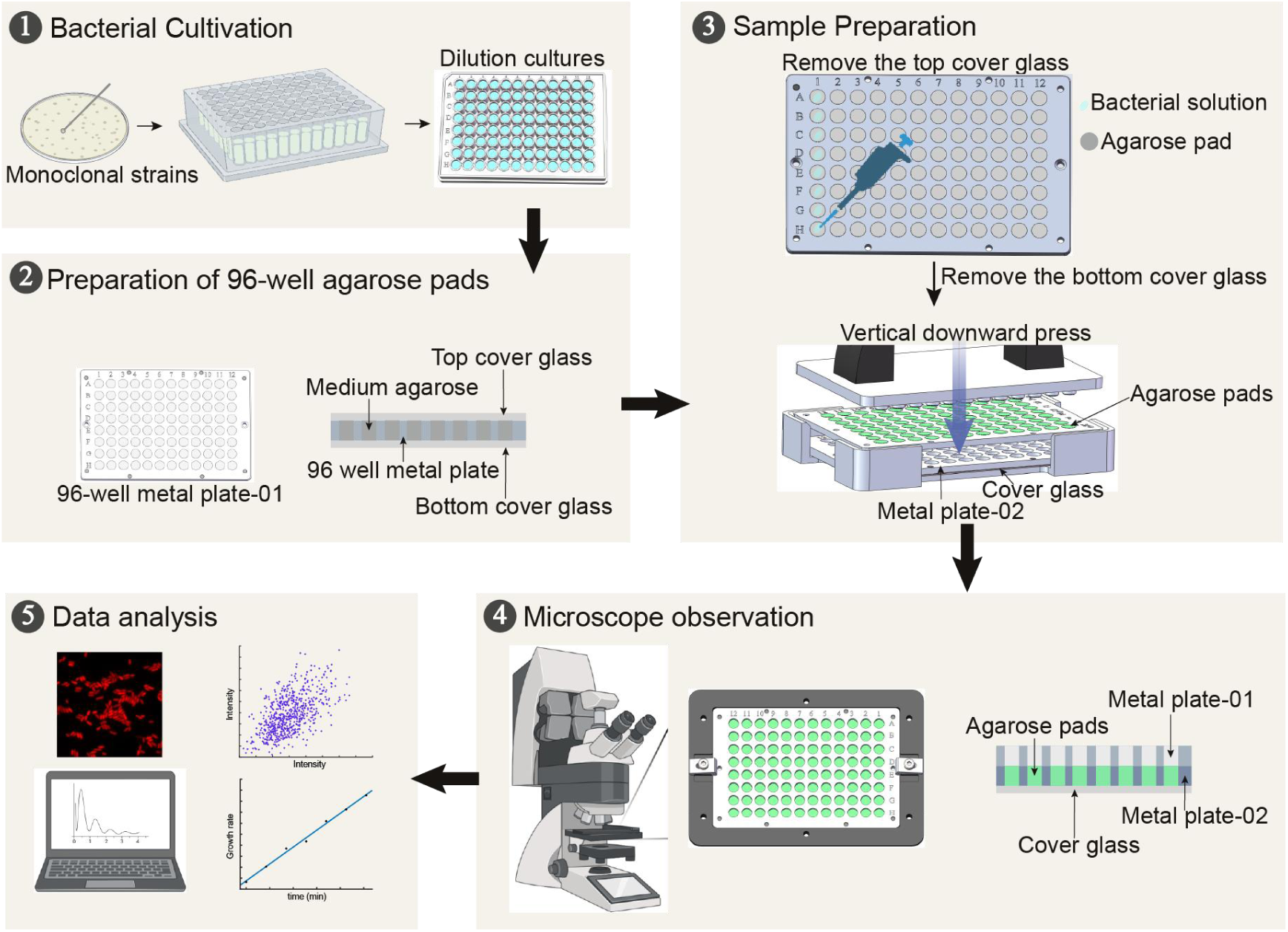
Sampling procedures for high-throughput microscopy-based platform for simultaneous analysis. (1) Bacterial cultivation. Bacterial strains recovered on LB agar were scraped from the plates and resuspended into the 96-deep-well plate for cultivation, and were further diluted into the 96-well plate for following sample preparation. (2) Preparation of 96-well agarose pads. The 96-well agarose pad plates were prepared by sandwiched between top and bottom cover glass in the first 96-well metal plate. (3) Sample preparation. Tipping the diluted bacterial solution onto 96-well agarose pad after removing the top slide glass, and sandwiching the bacteria between the agarose pad and cover glass with the downward press in vertically into the second metal plate after removing the bottom slide glass. (4) Microscope observation. The two metal plates harboring bacteria samples were anchored with magnetic force, then transferred onto the microscope for observation. (5) Data analysis. The acquired fluorescent images were processed and analyzed by using MATLAB with a self-written code.

### Screening of a natural promoter library

In a previous study, we constructed an dual-color reporter library consisting of 3,025 natural intrinsic promoters from *Pseudomonas aeruginosa*^25^. This library employed *sfGFP* as a quantifiable marker to measure the activity of each promoter, while the constitutive promoter J23102, fused to the gene of *CyOFP1* fluorescent protein, served as an internal control (Figure 2a). To evaluate the promoter activity with high throughput, we utilized our 96-well pad plate platform to acquire dual fluorescent images of bacteria and analyze the resulting images to determine the fluorescent intensity (Figure 2b). Initially, we screened a subset of 96 promoters from the initial library (Table S1) and evaluated the fluorescence of sfGFP and CyOFP1 for each reporter in both exponential and stationary growth phases in FAB medium (Figure 2c). For the constitutively expressed control gene, all strains exhibited a 2-fold higher fluorescence in CyOFP1 in stationary phase than exponential phase (Figure 2c), indicating that the growth conditions indeed influence protein expression, possibly due to changes in protein dilution time^26^. Regarding the sfGFP fluorescent reporter, we observed that a majority of promoters exhibited low activity, as indicated by sfGFP fluorescence levels below 50 a.u. (approximately 82%, 80 out of 96). To identify robust promoter candidates, we postulated that they should exhibit moderate to high activity, while maintaining stable expression levels under varying growth conditions. As such, we screened twelve strains that displayed high sfGFP fluorescence greater than 50 a.u., with fluorescence differences no greater than 40% between the exponential and stationary phases (as indicated by the arrows in Figure 2c).

**Figure 2.**
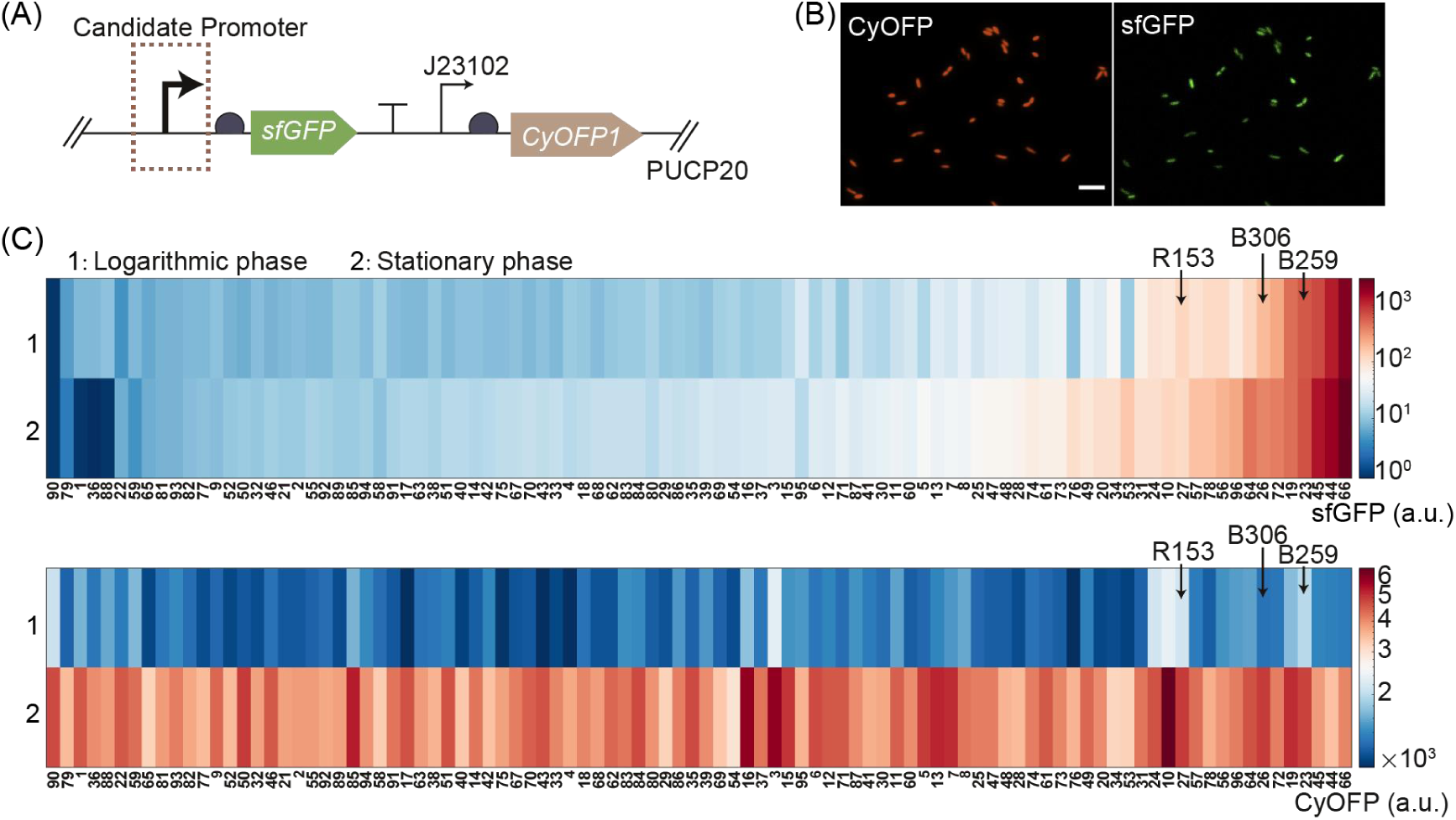
Screening of a natural promoter library. (A) In the promoter library, the sfGFP fluorescent protein served as a quantifiable marker, while CyOFP1 fluorescent protein served as an internal control by fused with a constitutive promoter J23102, in the vector PUCP20. (B) The representative fluorescent images in CyOFP1 and sfGFP channel. Here the scale bar is 5 μm. (C) Comparison of the sfGFP and CyOFP1 fluorescent intensity in logarithmic phase and stationary phase for each promoter from the promoter library.

We expanded our investigation to include five different types of growth conditions, which consisted of various culture media and stress conditions, such as rich medium, iron-deprivation, and antibiotic addition (See Methods for detail). To assess the impact of endoplasmid on bacterial growth, we employed a microplate reader to continuously monitor the growth curves of 12 strains harboring different promoter reporter plasmids across 8 distinct media conditions, as depicted in Figure S3. The observed overlap in the growth curves implies that the presence of endoplasmids, along with the different promoter reporter plasmids, did not significantly alter the growth dynamics of the bacteria under the tested conditions. We then measured the corresponding activity of the selected 12 promoters in *P. aeruginosa* under these conditions (Figure 3). In line with the results depicted in Figure 2c, the fluorescence intensity of CyOFP1 in all strains displayed considerable variability across different conditions (Figure 3b), with a high coefficient of variation (CV) of greater than 0.60 (CV: the ratio of the standard deviation to the mean).

**Figure 3.**
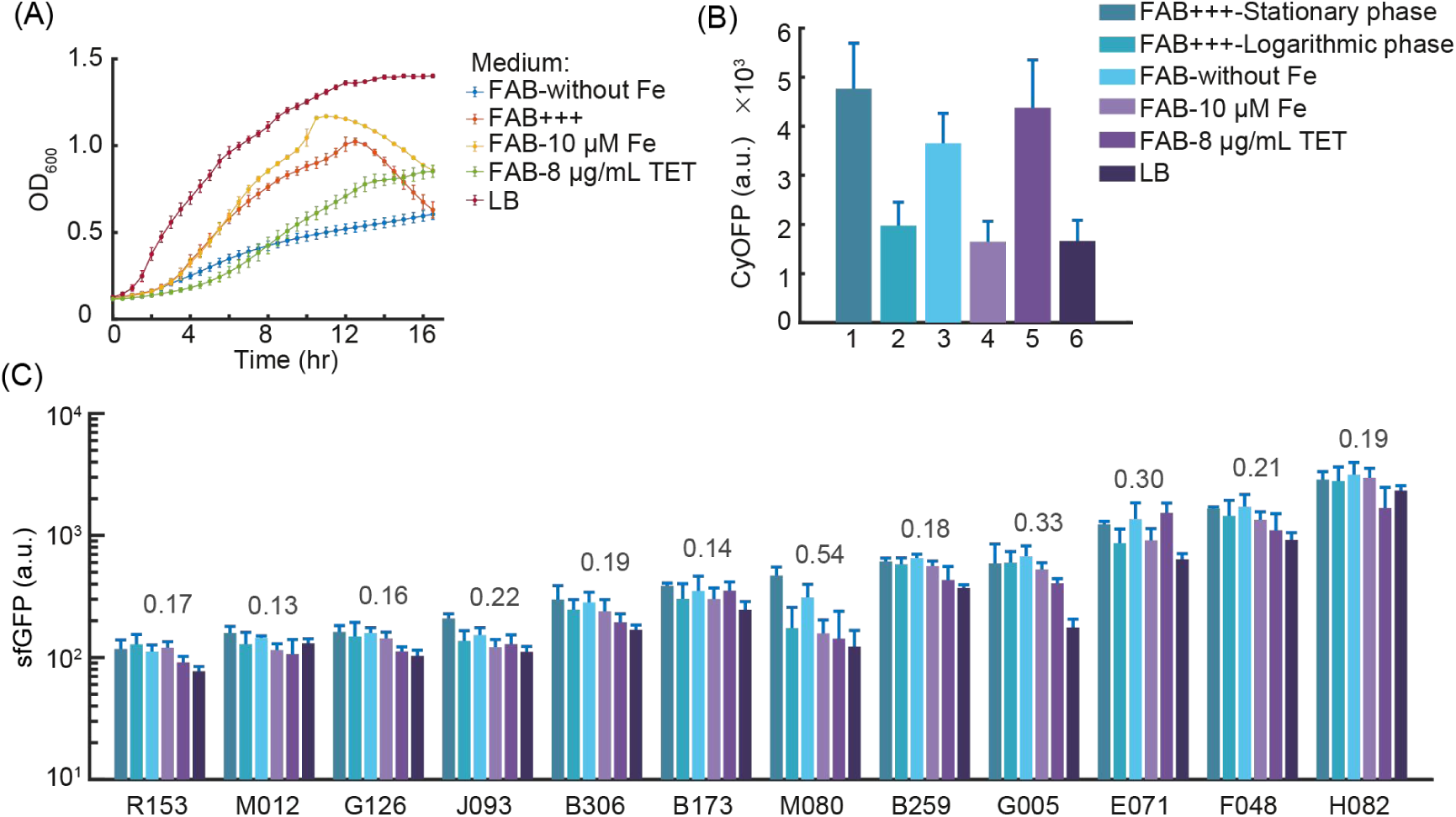
Characterization of selected promoters from the promoter library in 5 different types of growth conditions. (A) OD_600_ of the selected promoters were measured in 5 types of growth conditions, including FAB without FeCl_3_ (FAB-without Fe, marked with blue line), normal FAB media (FAB+++, marked with orange line), FAB with 10 μM iron (FAB-10 μM Fe, marked with yellow line), FAB with antibiotic of tetracycline (FAB-8 μg/mL TET, marked with green line), rich medium LB (LB, marked with brown line). (B) The average intensity of CyOFP1 of the selected promoters were measured in 6 different conditions, including normal FAB media in stationary phase (FAB+++-Stationary phase), normal FAB media in logarithmic phase (FAB+++-Logarithmic phase), FAB without FeCl_3_ (FAB-without Fe), FAB with 10 μM iron (FAB-10 μM Fe), FAB with antibiotic of tetracycline (FAB-8 μg/mL TET), rich medium LB (LB), respectively. (C) The average intensity of sfGFP of the selected 12 promoters (R153, M012, G126, J093, B306, B173, M080, B259, G005, E071, F048, H082) were measured in the same conditions as (b). Data are presented as mean ± SD (standard deviation, n = 3). The CV values for each set of data are shown in grey font. The coefficient of variation (CV) is calculated by the ratio of standard deviation to mean.

In contrast, all 12 measured promoters showed moderate to high activity across all conditions, with half of the promoters (7 of 12) exhibiting high conservation in expression levels across growth conditions in *P. aeruginosa*, as evidenced by a low CV of 0.2 for sfGFP fluorescence (Figure 3c).Together, by leveraging our 96-well pad plate platform for screening, we successfully identified 7 robust promoter candidates that exhibited minimal sensitivity to factors such as growth phase or culture medium composition in *P. aeruginosa*.

### Characterization of promoter activity at the single cell level

One advantage of microscopy-based screening is the ability to track temporal processes in single cells, and our platform is no exception. To illustrate this, we representatively performed single-cell analysis of the B173 promoter reporter strain under normal and antibiotic-added FAB medium conditions, as shown in Figure 4. In contrast to one-shot image scanning, we performed time-lapse imaging over a period of 2 hours with 17-minute intervals (Figure 4a) and made single-cell measurements of cell growth, such as growth rate, by fitting the time course data of cell area (Figure 4b). In all the experimental conditions we tested, *P. aeruginosa* grew the fastest in LB (Figure S4 and Figure 3a). We first conducted a quantitative characterization of the growth rate of *P. aeruginosa* in LB medium. As shown in Figure S5, we present accurate measurements of bacterial growth rate, indicating that the current image interval (17 min) employed in the *P. aeruginosa* strains meets the experimental requirements. We analyzed more than 10^3^ cells and found that the addition of antibiotics significantly decreased the mean cell growth rate (0.004 min^-1^ vs 0.002 min^-1^, p < 0.001. Figure 4c), despite the final antibiotic concentration being below the minimal inhibitory concentration (MIC). Furthermore, we generated scatter plots to visualize the relationship between growth rate and fluorescence intensity at the single cell level (Figure 4d-4f). The results presented in Figure 4e demonstrate that the sfGFP fluorescence intensity in the bacteria remained constant regardless of variations in the growth rate. While, as the growth rate decreased, the expression of CyOFP1 increased (Figure 4f, and Figure S4). The results demonstrated a negative correlation coefficient of −0.52 between the fluorescence intensity of constitutively expressed CyOFP1 and growth rate (Figure 4d), whereas the correlation coefficient between sfGFP fluorescence and growth rate was close to zero, indicating no relationship (Figure 4e and 4f). To further validate the robust function of the promoter at different growth phases, we also performed single-cell analysis of dual-color fluorescence at different culture times. The results revealed a downward trend in the growth rate of bacteria as the culture time increased, indicating that the presence of TET in the medium had a negative impact on bacterial growth. The histograms of the fluorescence intensity revealed that the fluorescence intensity of CyOFP driven by the constitutive promoter increased with culture time (0, 2, 4, and 8 h), whereas the fluorescence intensity of sfGFP driven by B173 promoter remained stable during cell growth (Figure 4g). Based on these single-cell data, we confirmed that the activity of the B173 promoter is independent on the environment stress.

**Figure 4.**
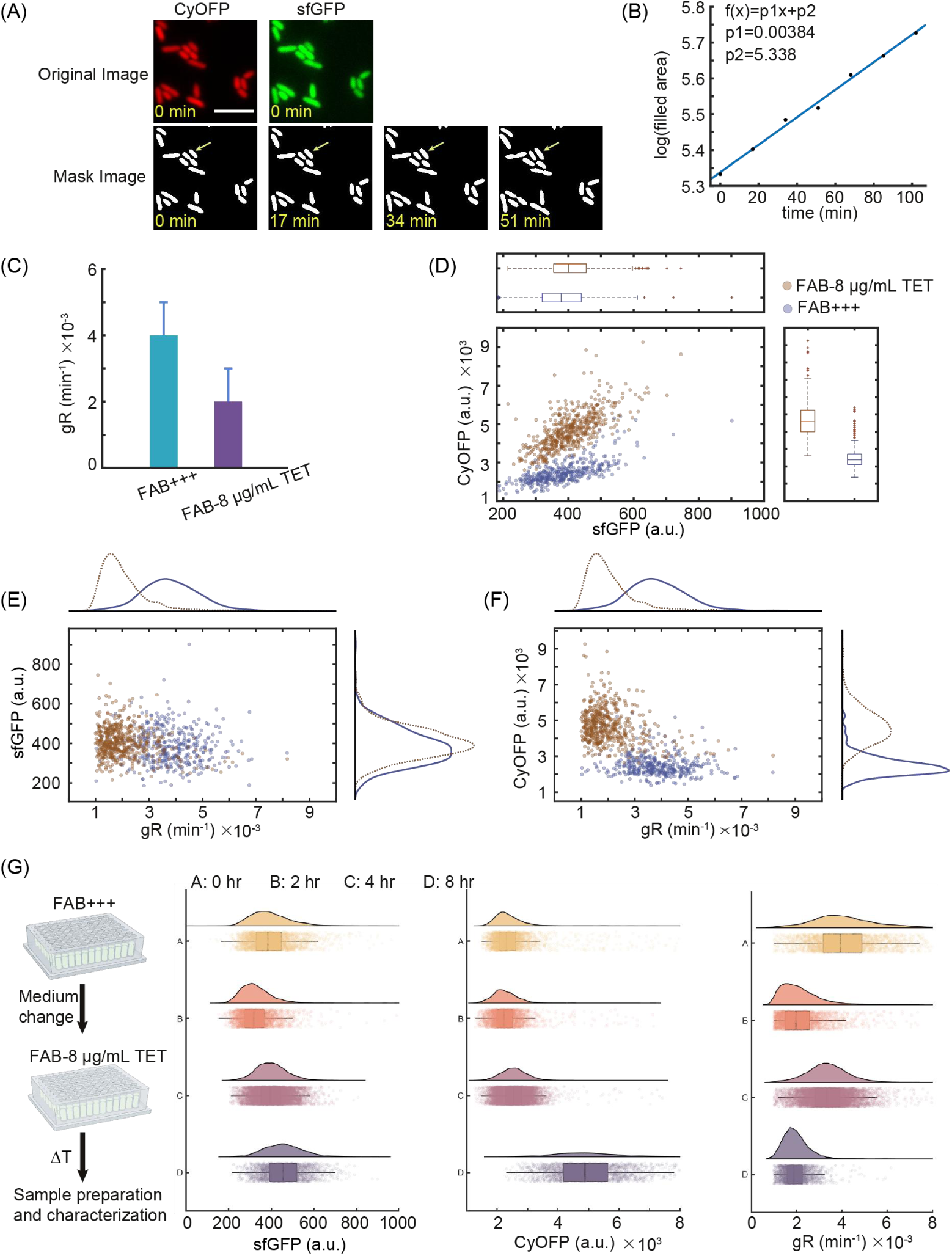
Characterizing the promoter activity of the selected promoter (B173) from the promoter library at single cell level. (A) The time-lapse images were performed over a period of 2 hours with 17-minute intervals with the mask images conducted from the original sfGFP and CyOFP1 images, where the yellow arrow indicated a tracked single cell. Here the scale bar is 5 μm. (B) Measuring the growth rate of single cell by fitting the time course data of cell area. (C) The mean cell growth rate was calculated by analyzing more than 10^3^ cells in FAB with antibiotic of tetracycline (FAB-8 μg/mL TET), compared with that in normal FAB media. Data are presented as mean ± SD (standard deviation, n >10^3^). Visualizing the relationship between the sfGFP and CyOFP1 intensity (D), between the growth rate and the sfGFP intensity (E), and between the growth rate and the CyOFP1 intensity (F) at the single cell level by generated scatter plots in FAB with antibiotic of tetracycline (FAB-8 μg/mL TET), compared with that in normal FAB media. (G) The single-cell analysis of dual-color fluorescence was performed at different culture times (0, 2, 4 and 8 hours) with the histograms of the sfGFP intensity, CyOFP1 intensity and growth rate, by changing the medium from normal FAB media to FAB with antibiotic of tetracycline.

### Stable protein expression arises from consistent transcriptional activity

Whereas protein expression involves the processes of transcription of a promoter and translation of information from a specific gene, we aimed to investigate whether stable expression of fluorescent proteins, driven by candidate robust promoters, depends on the ribosome binding site (RBS) or the coding sequence. We replaced the original RBS of four selected promoter reporters with three different constitutive prokaryotic RBSs of varying strengths and measured sfGFP expression in *P. aeruginosa* under the aforementioned five growth and stress conditions. We observed varying fluorescence intensity with different RBS strengths, but consistent fluorescence levels across the five different growth conditions for each RBS (coefficient of variation [CV] < 0.17 for all reporters; Figure 5a and 5c). Furthermore, we replaced the original *sfGFP* coding sequence with that of another fluorescent protein, CyOFP1, and still observed stable fluorescence levels across different growth conditions (Figure 5a and 5b). These results suggest that the stable protein expression driven by the screening promoter arises from its robust transcriptional activity, rather than post-transcriptional processes.

**Figure 5.**
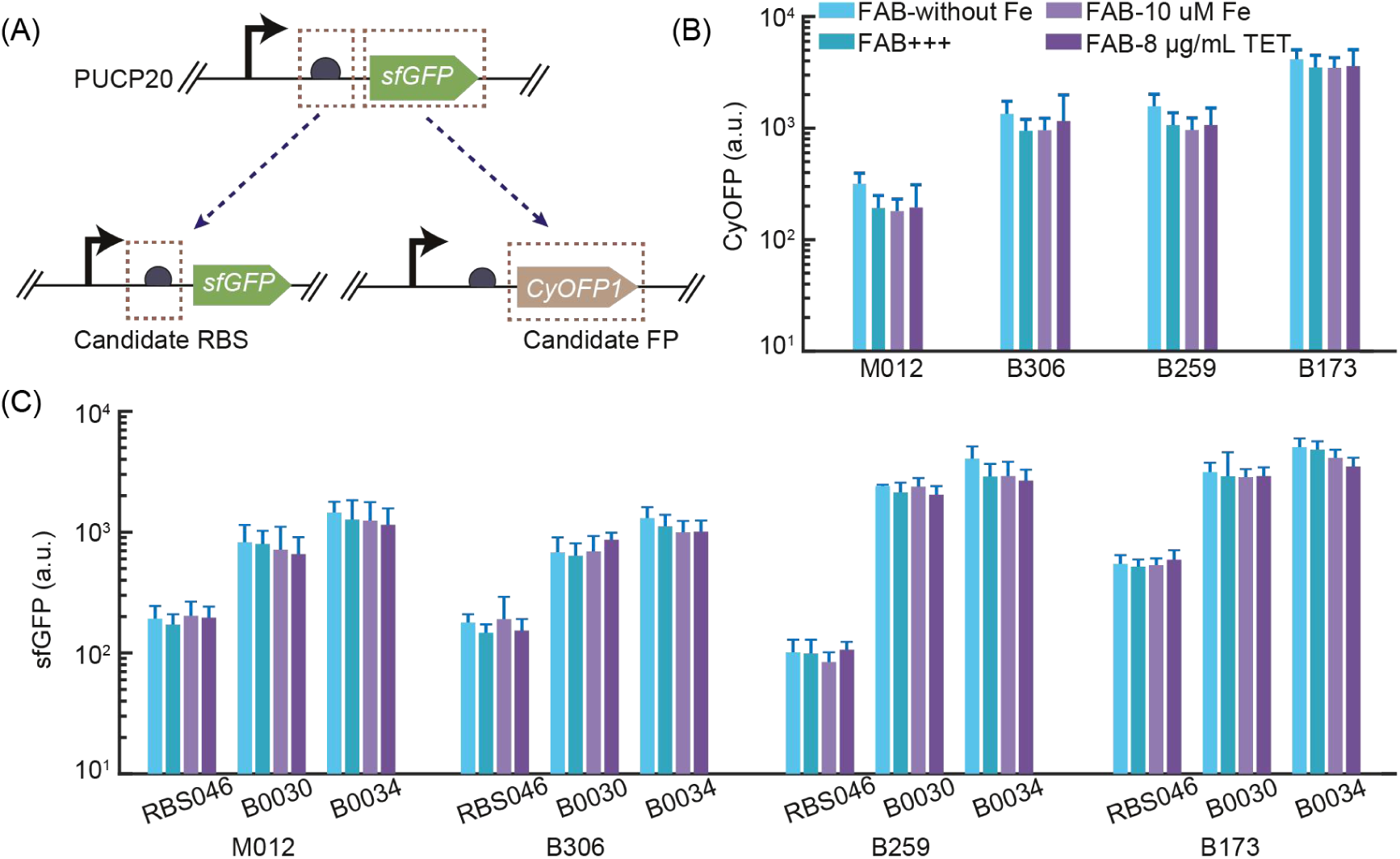
Characterization of the four robust promoters from the promoter library with replaced RBS or reporter fluorescence protein. (A) The schematic diagram of promoter with replaced RBS or reporter fluorescence protein (transfer to CyOFP1). (B) The CyOFP1 intensity of the four robust promoters (M012, B306, B259, B173) with the replaced reporter fluorescence protein (CyOFP1) were measured in 4 different conditions, including FAB without FeCl_3_ (FAB-without Fe), normal FAB media (FAB+++), FAB with 10 μM iron (FAB-10 μM Fe), and FAB with antibiotic of tetracycline (FAB-8 μg/mL TET). (C) The sfGFP intensity of the four robust promoters (M012, B306, B259, B173) with the replaced RBS (RBS046, B0030, B0034) were measured in 4 different conditions, including FAB without FeCl_3_ (FAB-without Fe), normal FAB media (FAB+++), FAB with 10 μM iron (FAB-10 μM Fe), and FAB with antibiotic of tetracycline (FAB-8 μg/mL TET). Data are presented as mean ± SD (standard deviation). The experiment was repeated three times.

### Identification of the core sequence responsible for stable transcriptional activity in candidate robust promoters

When faced with unknown promoters and unclear regulatory mechanisms of promoter robustness, one approach is to construct a series of promoter truncated strains to systematically identify the core sequence of the promoters. By systematically truncating the promoter sequence, it becomes possible to identify essential regions or motifs that contribute to the promoter’s activity. Due to the length of the initial promoter sequences being over 800 bp, redundancies may exist. We selected four screened promoters and progressively truncated them using different primer pairs (Figure 6a; see methods for details). This truncated promoter library was then used to identify the core sequence responsible for stable transcriptional activity. We screened the truncated promoter with the criterion of maintaining similar strength as the native promoter template while still exhibiting stable expression levels under varying conditions. Figure 6b-e shows the sfGFP expression levels of a partially truncated promoter library in *P. aeruginosa* under two conditions, and we ultimately identified the core sequences of the four selected promoters. These core sequences exhibited similar robustness but were much shorter, being less than 200 bp (Table S4).

**Figure 6.**
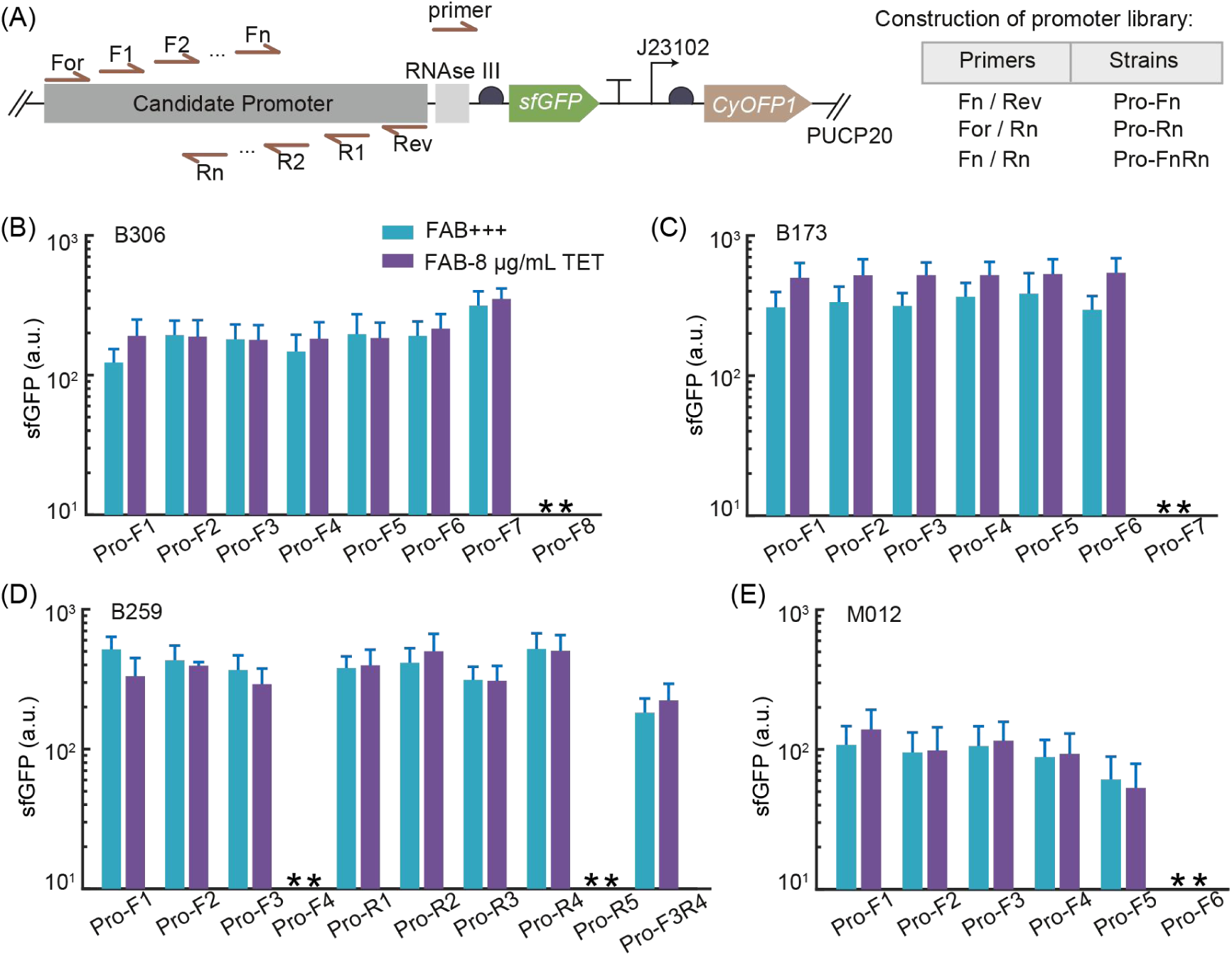
Identifying the core sequence responsible for stable transcriptional activity in candidate robust promoters. (A) The schematic diagram indicated the candidate promoters designed by truncating the promoters with different primers in vector PUCP20 and the construction of promoter library. (B-E) The sfGFP intensity of the candidate promoters (B306, B173, B259, M012, respectively) were measured in the normal FAB media (FAB+++), and FAB with antibiotic of tetracycline (FAB-8 μg/mL TET), where the truncated promoters referred to the promoter library in (A). Data are presented as mean ± SD (standard deviation). Experiments are replicated three times.

### Measuring the performance of candidate robust promoters in alternative host

We proceeded to utilize our established 96-well pad plate platform to conduct high-throughput quantification of the fluorescence output generated from each of the candidate robust promoters in a different recipient, *Pseudomonas putida* KT2440. We transformed the twelve promoter reporters from *P. aeruginosa* as depicted in Figure 2 and measured the sfGFP expression levels under five different growth and stress conditions (Figure 7). These promoters spanned nearly three orders of magnitude of fluorescence in *P. putida*, ranging from 40 to 4,000 a.u. Moreover, most of all promoters exhibited stable functions across all conditions, with a low coefficient of variation (CV <0.2) of fluorescence intensity among different conditions for each promoter (Figure 7). The results demonstrate the potential of these candidate promoters from *P. aeruginosa* to be expressed and applied in different host systems with flexibility.

**Figure 7.**
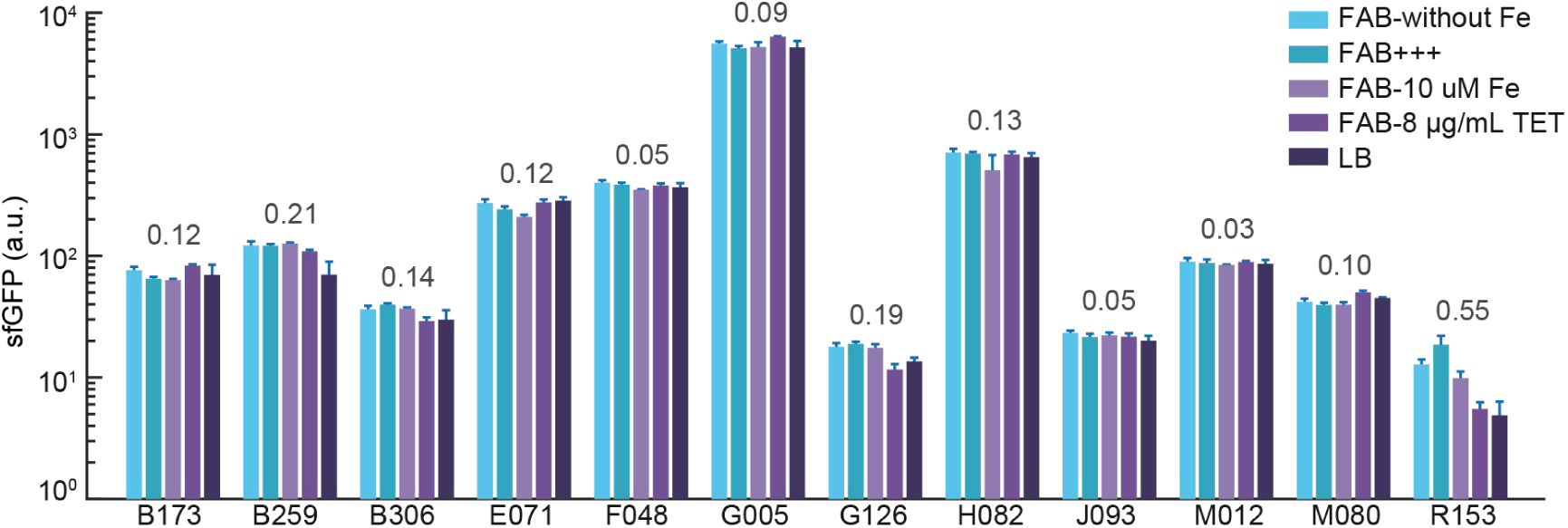
Measuring the promoter activity of candidate robust promoters in alternative host. The sfGFP intensity of the candidate robust promoters (B173, B259, B306, E071, F048, G005, G126, H082, J093, M012, M080, R153) were measured in different conditions (numbered 1 to 5) in alternative host *Pseudomonas putida* KT2440, including FAB without FeCl_3_ (FAB-without Fe), normal FAB media (FAB+++), FAB with 10 μM iron (FAB-10 μM Fe), FAB with antibiotic of tetracycline (FAB-8 μg/mL TET), rich medium LB (LB). The CV values for each set of data are shown in grey font. Data are presented as mean ± SD (standard deviation). Experiments are replicated three times.

## DISCUSSION

Current microscopy-based experiments can achieve high throughput in imaging billions of cells and enable long investigation times spanning several hours^19, 20^. However, they often face a major limitation in sample preparation, namely the inability to measure hundreds of samples in a high-throughput manner ^2, 4–6^. In this study, we have addressed this limitation by developing a novel high-throughput microscopy-based platform. This platform makes use of a high-throughput micro-sampling device that enables the simultaneous preparation of an 8×12 well agarose pads plate. This plate is then compatible with standard microscope object stages, allowing for the imaging screening of up to 96 independent strains or experimental conditions in a single experiment. By utilizing this platform, we were able to significantly increase the efficiency of our experiments, while also maintaining a high level of accuracy and precision in our measurements.

To highlight the flexibility, the platform was utilized to screen a library of natural promoters and identify robust promoter candidates that exhibited minimal sensitivity to factors such as growth phase or culture medium composition in *P. aeruginosa*. We then validate the robustness of these promoters in an alternative host, *P. putida* KT2440, and that all twelve candidate promoters retained stable functions across all tested conditions. These candidate promoters have the potential to be used in different host systems. Researching and applying robust promoters can enhance our understanding of gene regulation mechanisms and have important applications in synthetic biology, genetic engineering, and biomedical research.

Robust promoters represent a valuable tool for regulating gene expression in various biological applications, such as synthetic biology, metabolic engineering, and gene therapy. By providing more predictable and consistent gene expression levels under varying growth conditions, they can minimize the impact of environmental factors that could affect gene expression. This makes them particularly useful for designing and constructing synthetic gene circuits that require precise regulation of gene expression to achieve specific functions^27^. Moreover, robust promoters can enhance the production of useful biomolecules, such as therapeutic proteins or biofuels^28, 29^, by ensuring consistent and high-level expression of the target genes. The use of a promoter library can also expand promoter diversity, improving DNA assembly efficiencies for larger and more complex gene circuits, while maintaining their evolutionary stability^30, 31^. In this study, we attempted to elucidate the factors contributing to the robustness of the screened promoters by employing a diverse range of strategies (Table S6). However, despite our diligent endeavors, a definitive mechanism explaining this robustness remains elusive. Our dedication to unraveling the underlying mechanisms will persist, and we aim to continue exploring and probing this intriguing aspect of promoter behavior to gain a more comprehensive understanding. Overall, robust promoters play a significant role in maintaining stable gene expression, reliable signal transduction, consistent gene expression, and mutation resistance. The identification and use of robust promoters has the potential to revolutionize the field of synthetic biology and enable the development of more efficient and reliable biological systems for a wide range of applications.

In light of the stable activity exhibited by these promoters, it is likely that they are regulated by different sigma factors that mediate non-equivalent levels of transcription in response to diverse milieu^27, 32, 33^. Alternatively, they may be subject to negative feedback coupling with growth rate or burden^34–36^, or other unknown mechanisms. Although unexplored in this study, future investigations could focus on unraveling the underlying mechanisms responsible for the stable gene expression of these candidate promoters.

Another key advantage of our established platform is that it allowed for the tracking of temporal processes in single cells, which enabled us to quantify cell features related to time series such as growth rate (Figure 4). This capability also allowed for analysis of gene expression dynamics, making high-throughput screening of biosensors possible^37^. For instance, using our platform, we could screen fluorescent biosensors by monitoring the real-time fluorescence changes in response to a specific messenger using microscopy in intact model organisms. With a capacity to screen 96 variants in high throughput, this approach could significantly increase the efficiency and speed of biosensor screening.

The platform utilized in this study has some limitations. While the concentration of agarose can be adjusted to allow for twitching motility when cells are grown under agarose pads, swimming motility cannot be assessed, which limits screening of properties related to swimming. Additionally, although small-molecule inducers or chemical perturbations can be easily added to the top of the pad while the bacteria are growing, their removal is difficult. However, the transparency of the agarose pads allows for the integration of optogenetic devices and genetic circuits to change or control input signals with light^38^. Another limitation of the platform is that while it allows for the tracking of temporal processes, tracking cell lineages for arbitrarily long timescales is not practical due to the limited nutrients and restricted space. As a solution, custom devices could be designed to continuously supply fresh medium to cells beyond the agarose, similar to the mother machine^39^.

Overall, the platform described in this article represents an advance in microscopy-based screening approaches and has the potential to facilitate the identification of genetic elements with temporal or spatial properties such as biosensor kinetics, gene expression dynamics. We demonstrated the platform’s flexibility by using it to screen and quantify robust promoters. The platform has the potential to be adapted for use with other microbial organisms, with some necessary adjustments, such as modifying the composition of agarose pads, to provide a powerful tool for high-throughput genetic analysis. We anticipate that this platform’s versatility and ability to enable quantification in high throughput will lead to its broad adoption in various fields, including biomedical sciences, bioengineering, and synthetic biology.

## MATERIALS AND METHODS

### Bacterial strains and growth conditions

The bacterial strains used in this study include *Pseudomonas aeruginosa* strain PAO1 and *Pseudomonas putida* KT2440. The screened promoters of initial library are based on the dual-color reporter library of 3025 natural intrinsic promoters from *Pseudomonas aeruginosa* from previous study. The screened promoters and the corresponding gene name are listed in Tables S1, and the detailed sequences of 12 analyzed promoters are enclosed in Tables S2. All bacterial culture experiments were conducted at 37°C. Strains carrying reporter plasmids were grown in the classical LB broth (10 g/L NaCl, 10 g/L tryptone, 5 g/L yeast extract) supplemented with 30 μg/ mL gentamicin overnight, following with a dilution into FAB medium with 30 mM glutamate as a carbon source under aerobic conditions, with the composition as follows per liter: (NH_4_)_2_SO_4_, 2 g/L; Na_2_HPO_4_ · 12H_2_O, 12.02 g/L; KH_2_PO_4_, 3 g/L; NaCl, 3 g/L; MgCl_2_, 93 g/mL; CaCl_2_·2H_2_O, 14 g/mL; FeCl_3_, 1 μM; and trace metals solution (CaSO_4_·2H_2_O, 200 mg/L; MnSO_4_·7H2O, 200 mg/L; CuSO_4_·5H_2_O, 20 mg/L; ZnSO_4_ ·7H_2_O, 20 mg/L; CoSO_4_ ·7H_2_O, 10 mg/L; NaMoO_4_ ·H_2_O, 10 mg/L; H_3_BO_3_, 5 mg/L) 1 mL/L. The FAB media with 10 μM FeCl_3_ or without Fe or with 8 μg/mL tetracycline were used when tested in different culture conditions. The shaker for cultivating bacteria was set at 800 rpm in a deep 96-well plate.

### Construction of plasmids and strains

All plasmids and strains were listed in Table S3. All plasmids were constructed using basic molecular cloning techniques and Gibson assembly. All plasmids were sequenced and then transformed into PAO1 or KT2440 with standard protocols^40, 41^. The original plasmids used for calibrating the robustness of promoter contain two modules: *CyOFP1* driven by J23102 was used as an internal control, and *sfGFP* driven by fixed promoter was the reporter. To investigate whether stable expression of sfGFP, driven by candidate robust promoters (e.g. B173), depends on the RBS or the coding sequence, we replaced the RBS and the coding sequence sfGFP of candidate promoter in template plasmid of B173-sfGFP-J23012-CyOFP1-PUCP20 with RBS046/B0030/B0034 and CyOFP respectively. B173-CyOFP1-PUCP20 was constructed by replacing *sfGFP* in B173-sfGFP-J23012-CyOFP1-PUCP20 with *CyOFP1* via Gibson assembly. The vector fragment B173-PUCP20 without J23102-CyOFP1 was obtained by PCR, and then connected with sfGFP fragment to construct the plasmid B173-CyOFP1-PUCP20. PCR fragments sfGFP together with various RBS was ligated with linearized vector fragment B173-J23012-CyOFP1-PUCP20 to generate plasmid B173-RBS046/B0030/B0034-sfGFP-J23012-CyOFP1-PUCP20.

In order to identify the core sequences responsible for stable transcriptional activity among candidate robust promoters, we constructed serial truncations of the promoter into dual-color reporter vector and then validated the robustness of the truncated promoters in PAO1 cells. The schematic diagram of the construction of truncated promoter libraries is shown in Figure 6a. Taking the truncations of B173 promoter as an example, the detailed experimental process is as follows: A series of PCR primers were designed to amplify different truncations of the candidate robust promoters; the PCR fragments were cloned into reporter plasmid PUCP20 and placed upstream of sfGFP; the resultant plasmids were finally electroporated into PAO1 to generate the truncated promoter library.

### Preparing 96-well agarose pads

The 96-well agarose pad plates were prepared by sandwiched between two (top and bottom) cover glass in the 96-well metal plates. These plates have been electroplated on the surface to enhance their durability and resistance to corrosion, making them suitable for reuse. After each experiment, the device is subjected to two separate cleaning steps. Firstly, the device is immersed in 75% alcohol and subjected to ultrasonic cleaning for 10 minutes. This helps to remove any residual substances or contaminants. Afterward, the device is cleaned with water using the same ultrasonic cleaning process. Once the cleaning process is complete, the device is dried and ready for reuse. To ensure accurate and high-quality imaging results, fresh slides are necessary for each experiment. The cover glass was ultrasonic cleaned with 75% ethanol and Millipore water respectively for 10 minutes. The detailed process of preparing 96-well agarose pads is shown in Figure S1 and S2. Firstly, we glued a piece of cover glass on the bottom surface of sheet metal −01, and then flipped the metal plate so that the top surface was facing up. Secondly, the plastic choke plate was placed on the top surface of metal plate that was locked in a waste tank. Third, bacterial culture medium containing 1% agarose was prepared, microwaved to melt completely, and then poured into the 96 holes in the metal plate. Next, we removed the choke plate and taken out a clean cover glass. Then, let one side of the cover glass contact with the top surface of the agarose, and attach the cover glass tightly to the top surface of the metal plate to ensure that there are no air bubbles between the cover glass and metal plate. Finally, the device is left at room temperature until the agarose in the holes is completely solidified.

### Sampling procedures for high-throughput microscopy-based platform

Bacterial strains were recovered from −80 ° C fridge on LB agar (1.5% w.t.) plates with 30 μg/mL gentamicin overnight at 37°C. Bacteria were scraped from the plates and resuspended in 1 mL of FAB media (96-deep well plate) with 30 μ g/mL gentamicin and 1 μ M FeCl_3_. Strains were cultured overnight at 37 °C, and then diluted 100 times into 1 mL specific medium in 96-deep well plates for 6-hour cultivation. To prevent the formation of bacterial clusters during the experiment, it is crucial to address the issue of high initial bacterial density in the liquid. Hence, it is essential to dilute the initial bacterial solution to a suitable concentration. The bacteria cultures from 96-deep well plates were further diluted with fresh media before sampling onto the 96-well agarose pad plate. The diluted bacterial solution is added to the samples using the 8-Channel Pipettes, and the entire process takes less than 5 minutes. If any holes are missed during the initial addition, and additional drops of the bacterial solution can be added to those specific holes to ensure proper coverage. Tipping the diluted bacterial solution onto 96-well agarose pad after removing the top slide glass, and sandwiching the bacteria between the agarose pad and cover glass with the downward press in vertically by matching the two metal plates after removing the bottom slide glass. Then transfer the 96-well plates into the microscope for further observation and data analysis.

### Fluorescent image acquisition and data analysis

All qualitative images presented are representative of at least three independent duplicate experiments. Fluorescent images of sfGFP and CyOFP1 were acquired simultaneously by using two Zyla 4.2 Scmos cameras (Andor) with a fluorescence microscope (IX-71, Olympus) equipped with a 100x oil objective. The fluorescence of both sfGFP and CyOFP1 was excited at 488 nm using a solid-state light source (Lumencor Spectra X), and collected with emission filters of 520/28 nm and 583/22 nm, respectively. The sample arrangement and corresponding strains name are shown in Table S5. The microscope program is set to load 96 fields at once, each field being an independent strain or culture condition. Four pictures were collected in each field, and the number of bacteria in each picture was nearly 500. It takes about 15 minutes to complete the image acquisition at one time point. In order to characterize the growth rate of bacteria, we need to collect images at least 5 time points with a time interval of 17 minutes.

Image processing was conducted by using MATLAB with a self-written code. sfGFP and CyOFP1 images were aligned using a built-in function (imwarp) and CyOFP1 images were used for bacterial cell identification. The single-cell analysis of dual-color fluorescence was conducted with specific MATLAB code, and scatter plots were generated for visualization with MATLAB. All experiments data are expressed as the mean ± SD.

## ACKNOWLEDGMENTS

We would like to thank all the lab members of Shenzhen Synthetic Biology Infrastructure for helpful discussions. This work was supported by the National Key Research and Development Program of China (Grant No. 2020YFA0906900), the National Natural Science Foundation of China (Grant No. 32000061 to RR.Z, 32101177 to YJ.H).

## AUTHOR CONTRIBUTIONS

RR.Z and YJ.H contributed equally to this work. F.J., S.Y, and RR.Z. designed the experiments. RR.Z. and YJ.H performed experiments and analyzed data. M.L and B.L helped to construct bacterial strains. AG.X. helped to obtain microscope images. L.W and Y.L helped designed and fabricated the device of high-throughput microscopy-based platform. RR.Z., YJ.H and S.Y contributed jointly to data interpretation and manuscript preparation. All authors reviewed the manuscript.

## DATA AVAILABILITY

All data used during the study appear in the submitted article.

## ETHICS STATEMENT

This article does not contain any studies with human participants or animals performed by any of the authors.

## REFERENCES

1. St Johnston D. The art and design of genetic screens: Drosophila melanogaster. Nat Rev Genet. 2002, 3 (3), 176–188.

2. Boutros M, Heigwer F, Laufer C. Microscopy-Based High-Content Screening. Cell. 2015, 163 (6), 1314–1325.

3. Benfey PN, Mitchell-Olds T. From genotype to phenotype: systems biology meets natural variation. Science. 2008, 320 (5875), 495–497.

4. Pepperkok R, Ellenberg J. High-throughput fluorescence microscopy for systems biology. Nat. Rev Mol Cell Biol. 2006, 7 (9), 690–696.

5. Zanella F, Lorens JB, Link W. High content screening: seeing is believing. Trends Biotechnol. 2010, 28 (5), 237–245.

6. Young JW, Locke JCW, Altinok A, Rosenfeld N, Bacarian T, Swain PS, Mjolsness E, Elowitz MB. Measuring single-cell gene expression dynamics in bacteria using fluorescence time-lapse microscopy. Nat. Protoc. 2012, 7 (1), 80–88.

7. Wang YH, Wei KY, Smolke CD. Synthetic Biology: Advancing the Design of Diverse Genetic Systems. Annu Rev Chem Biomol Eng. 2013, 4 (1), 69–102.

8. Mayer TU, Kapoor TM, Haggarty SJ, King RW, Schreiber SL, Mitchison TJ. Small Molecule Inhibitor of Mitotic Spindle Bipolarity Identified in a Phenotype-Based Screen. Science 1999, 286 (5441), 971–974.

9. Binan L, Mazzaferri J, Choquet K, Lorenzo LE, Wang YC, Affar EB, et al. Live single-cell laser tag. Nat. Commun. 2016, 7 (1), 11636.

10. Kuo CT, Thompson AM, Gallina ME, Ye F, Johnson ES, Sun W, et al. Optical painting and fluorescence activated sorting of single adherent cells labelled with photoswitchable Pdots. Nat Commun. 2016, 7 (1), 11468.

11. Chien MP, Werley CA, Farhi SL, Cohen AE. Photostick: a method for selective isolation of target cells from culture. Chem Sci. 2015, 6 (3), 1701–1705.

12. Chen KH, Boettiger AN, Moffitt JR, Wang S, Zhuang X. Spatially resolved, highly multiplexed RNA profiling in single cells. Science 2015, 348 (6233), aaa6090.

13. Moffitt JR, Hao J, Wang G, Chen KH, Babcock HP, Zhuang X. High-throughput single-cell gene-expression profiling with multiplexed error-robust fluorescence in situ hybridization. Proc Natl Acad Sci USA. 2016, 113 (39), 11046–11051.

14. Emanuel G, Moffitt JR, Zhuang X. High-throughput, image-based screening of pooled genetic-variant libraries. Nat Methods. 2017, 14 (12), 1159–1162.

15. Eng CHL, Lawson M, Zhu Q, Dries R, Koulena N, Takei Y, et al. Transcriptome-scale super-resolved imaging in tissues by RNA seqFISH+. Nature. 2019, 568 (7751), 235–239.

16. Feldman D, Singh A, Schmid-Burgk JL, Carlson RJ, Mezger A, Garrity AJ, et al. Optical Pooled Screens in Human Cells. Cell 2019, 179 (3), 787–799.e17.

17. Lee JH, Daugharthy ER, Scheiman J, Kalhor R, Ferrante TC, Terry R, et al. Fluorescent in situ sequencing (FISSEQ) of RNA for gene expression profiling in intact cells and tissues. Nat Protoc. 2015, 10 (3), 442–458.

18. Wang C, Lu T, Emanuel G, Babcock HP, Zhuang X. Imaging-based pooled CRISPR screening reveals regulators of lncRNA localization. Proc Natl Acad Sci USA. 2019, 116 (22), 10842–10851.

19. Lee J, Liu Z, Suzuki Peter H, Ahrens John F, Lai S, Lu X, et al. Versatile phenotype-activated cell sorting. Sci Adv. 2020, 6 (43), eabb7438.

20. Hasle N, Cooke A, Srivatsan S, Huang H, Stephany JJ, Krieger Z, et al. High-throughput, microscope-based sorting to dissect cellular heterogeneity. Mol Syst Biol. 2020, 16 (6), e9442.

21. Patterson GH, Lippincott-Schwartz J. A Photoactivatable GFP for Selective Photolabeling of Proteins and Cells. Science 2002, 297 (5588), 1873–1877.

22. Medaglia C, Giladi A, Stoler-Barak L, De Giovanni M, Salame TM, Biram A, et al. Spatial reconstruction of immune niches by combining photoactivatable reporters and scRNA-seq. Science. 2017, 358 (6370), 1622–1626.

23. Rimon N, Schuldiner M. Getting the whole picture: combining throughput with content in microscopy. J Cell Sci. 2011, 124 (22), 3743–3751.

24. Yang S, Cheng X, Jin Z, Xia A, Ni L, Zhang R, et al. Differential Production of Psl in Planktonic Cells Leads to Two Distinctive Attachment Phenotypes in Pseudomonas aeruginosa. Appl Environ Microbiol. 2018, 84 (14), e00700–18.

25. Chen W, Zhang J, Li F, Wang C, Zhang Y, Xia A, et al. Genome-Wide Analysis of Gene Expression Noise Brought About by Transcriptional Regulation in Pseudomonas aeruginosa. mSystems. 2022, 0 (0), e00963–22.

26. Taniguchi Y, Choi P J, Li GW, Chen H, Babu M, Hearn J, et al. Quantifying E. coli proteome and transcriptome with single-molecule sensitivity in single cells. Science. 2010, 329 (5991), 533–8.

27. Han L, Chen Q, Lin Q, Cheng J, Zhou L, Liu Z, et al. Realization of Robust and Precise Regulation of Gene Expression by Multiple Sigma Recognizable Artificial Promoters. Front Bioeng Biotech. 2020, 8.

28. Song Y, Nikoloff JM, Fu G, Chen J, Li Q, Xie N, et al. Promoter Screening from Bacillus subtilis in Various Conditions Hunting for Synthetic Biology and Industrial Applications. PLOS ONE 2016, 11 (7), e0158447.

29. Liu Y, Yap SA, Koh CMJ, Ji L. Developing a set of strong intronic promoters for robust metabolic engineering in oleaginous Rhodotorula (Rhodosporidium) yeast species. Microb Cell Fact. 2016, 15 (1), 200.

30. Sleight SC, Bartley BA, Lieviant JA, Sauro HM. Designing and engineering evolutionary robust genetic circuits. J Biol Eng. 2010, 4 (1), 12.

31. Torella JP, Lienert F, Boehm CR, Chen JH, Way JC, Silver PA. Unique nucleotide sequence–guided assembly of repetitive DNA parts for synthetic biology applications. Nat Protoc. 2014, 9 (9), 2075–2089.

32. Nicolas P, Mäder U, Dervyn E, Rochat T, Leduc A, Pigeonneau N, et al. Condition-Dependent Transcriptome Reveals High-Level Regulatory Architecture in Bacillus subtilis. Science. 2012, 335 (6072), 1103–1106.

33. Feklístov A, Sharon BD, Darst SA, Gross CA. Bacterial Sigma Factors: A Historical, Structural, and Genomic Perspective. Annu Rev Microbiol. 2014, 68 (1), 357–376.

34. Segall-Shapiro TH, Sontag ED, Voigt CA. Engineered promoters enable constant gene expression at any copy number in bacteria. Nat Biotechnol. 2018, 36 (4), 352–358.

35. Ceroni F, Boo A, Furini S, Gorochowski TE, Borkowski O, Ladak YN, et al. Burden-driven feedback control of gene expression. Nat Methods. 2018, 15 (5), 387–393.

36. Joshi SHN, Yong C, Gyorgy A. Inducible plasmid copy number control for synthetic biology in commonly used E. coli strains. Nat Commun. 2022, 13 (1), 6691.

37. Wang L, Wu C, Peng W, Zhou Z, Zeng J, Li X, et al. A high-performance genetically encoded fluorescent indicator for in vivo cAMP imaging. Nat. Commun. 2022, 13 (1), 5363.

38. Fu S, Zhang R, Gao Y, Xiong J, Li Y, Pu L, et al. Programming the lifestyles of engineered bacteria for cancer therapy. Natl Sci Rev. 2023, nwad031.

39. Wang P, Robert L, Pelletier J, Dang WL, Taddei F, Wright A, et al. Robust Growth of Escherichia coli. Curr Biol. 2010, 20 (12), 1099–1103.

40. Choi KH, Schweizer HP. mini-Tn 7 insertion in bacteria with single att Tn 7 sites: example Pseudomonas aeruginosa. Nat Protoc. 2006, 1 (1), 153–161.

41. Hoang TT, Kutchma AJ, Becher A, Schweizer HP. Integration-proficient plasmids for Pseudomonas aeruginosa: site-specific integration and use for engineering of reporter and expression strains. Plasmid. 2000, 43 (1), 59–72.

