## Supplementary material for "High-throughput, microscopy-based screening, and quantification of genetic elements": SI

**Affiliations:**
[1] CAS Key Laboratory of Quantitative Engineering Biology, Shenzhen Institute of Synthetic Biology, Shenzhen Institutes of Advanced Technology, Chinese Academy of Sciences, Shenzhen 518055, China;

[2] Shenzhen Synthetic Biology Infrastructure, Shenzhen Institute of Synthetic Biology, Shenzhen Institutes of Advanced Technology, Chinese Academy of Sciences, Shenzhen 518055, China;

[3]Chengdu Documentation and Information Center, Chinese Academy of Sciences, Chengdu 610299, China;

*To whom correspondence should be addressed.


[+]RR.Z and YJ.H contributed equally to this work.

#### Contents

| | |
|---|---|
| **Supplementary Figures 1** | **2** |
| **Supplementary Table 1-3** | **3--6** |
| **References** | **6** |

### Figure S1:

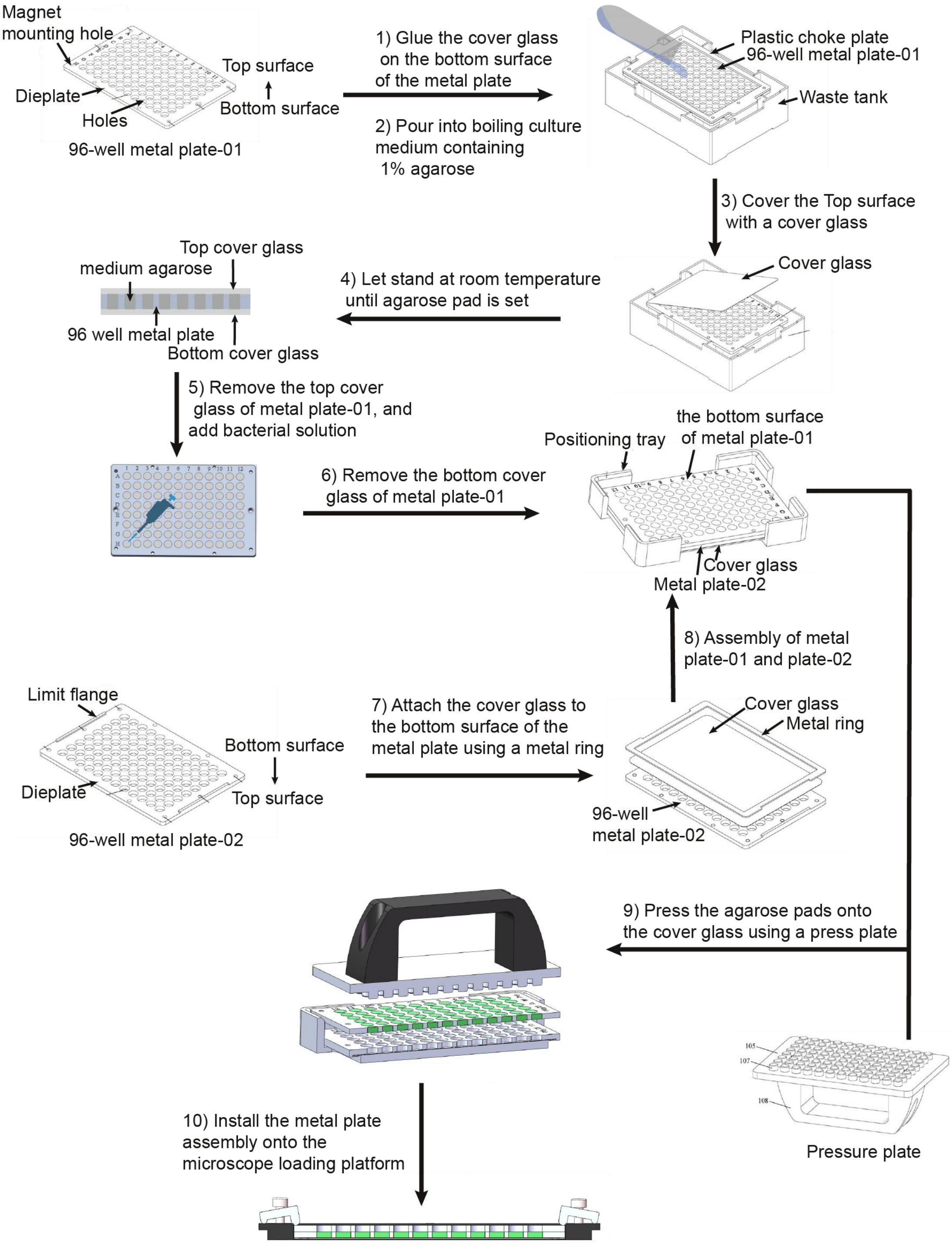


**Figure S1. The detailed procedures of sample preparation for high-throughput microscopy-based platform.** (1) The 96-well metal plate-01 consists of 96 holes in the die plate and a row of magnet mounting holes. Glue the cover glass on the bottom surface of the metal plate-01, then anchor the 96-well metal plate-01 into the plastic choke plate and locked in a waste tank, and (2) pour into boiling culture medium containing 1% (w/v) agarose. (3)

Cover the top surface with a piece of cover glass immediately. (4) Let stand at room temperature until agarose pad is set. (5) Removing the top cover glass of metal plate-01 and add bacterial solution via pipetting lightly. (6) Remove the bottom cover glass of the 96-well metal plate-01. (7) Attach the cover glass to the bottom surface of the 96-well metal plate-02 using a metal ring. (8) Assembly of the metal plate-01 (reverse vertically) and metal plate-02, and anchor them into the positioning tray. (9) Press the agarose pads onto the cover glass using a press plate. (10) Install the metal plates assembly onto the microscope loading platform for further acquisition of fluorescent images.

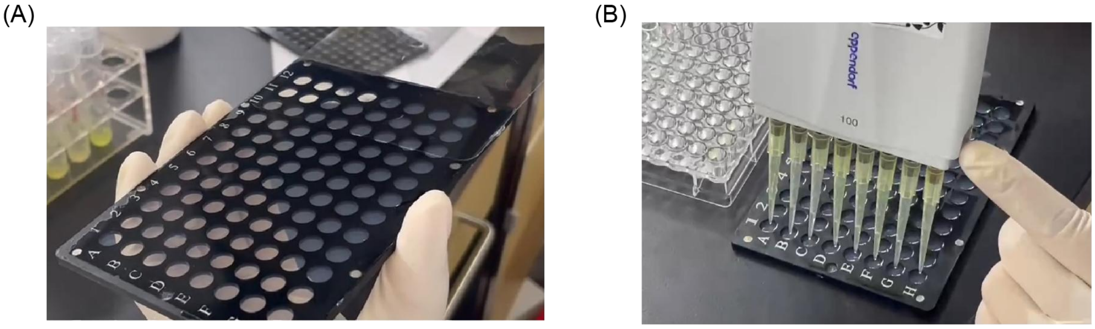

**Figure S2. Physical pictures of the microscopy-based platform.** (A) Process demonstration for making 96-well agar by removing the surface slide on the metal plate. (B) Application of bacteriological liquid to the agar surface using the 8-Channel Pipettes.

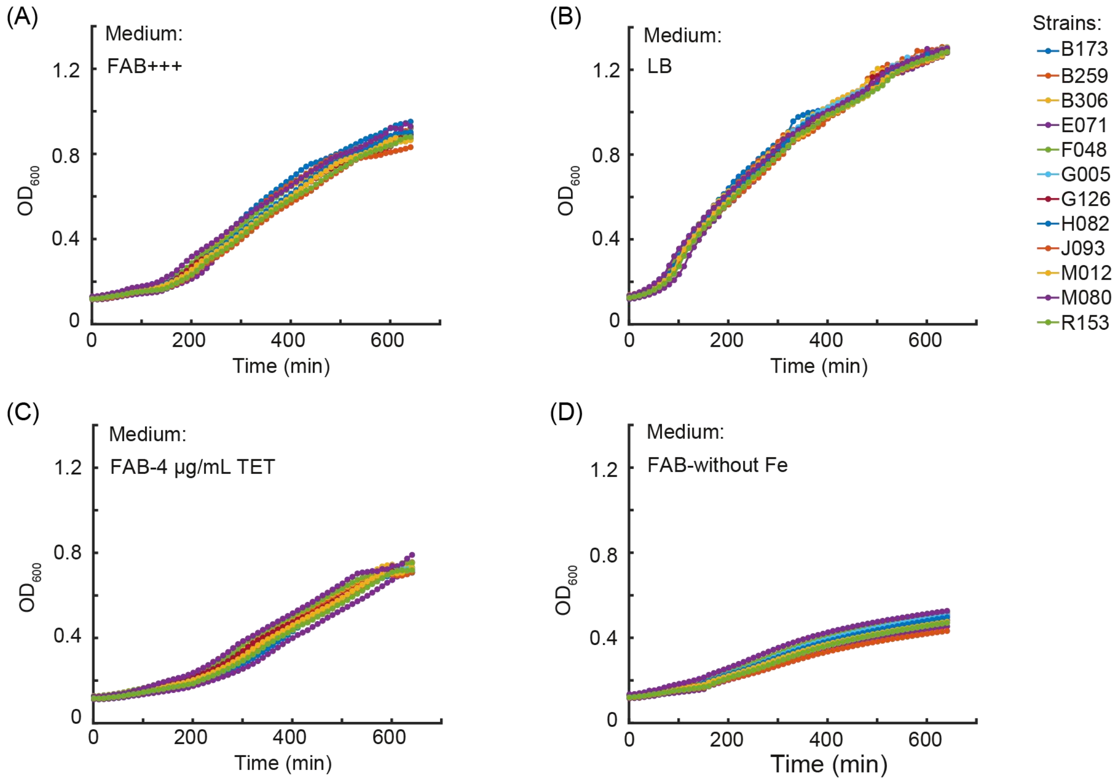


**Figure S3. The growth curves of 12 strains containing different plasmids were examined under various culture conditions.** A total volume of 100 μL medium was added to each well of a 96-well plate, and the logarithmic phase bacterial cultures were inoculated into different media. The growth of the strains was monitored under four distinct medium conditions (A) FAB+++, (B) LB, (C) FAB-4 μg/mL TET, (D) FAB-without Fe.

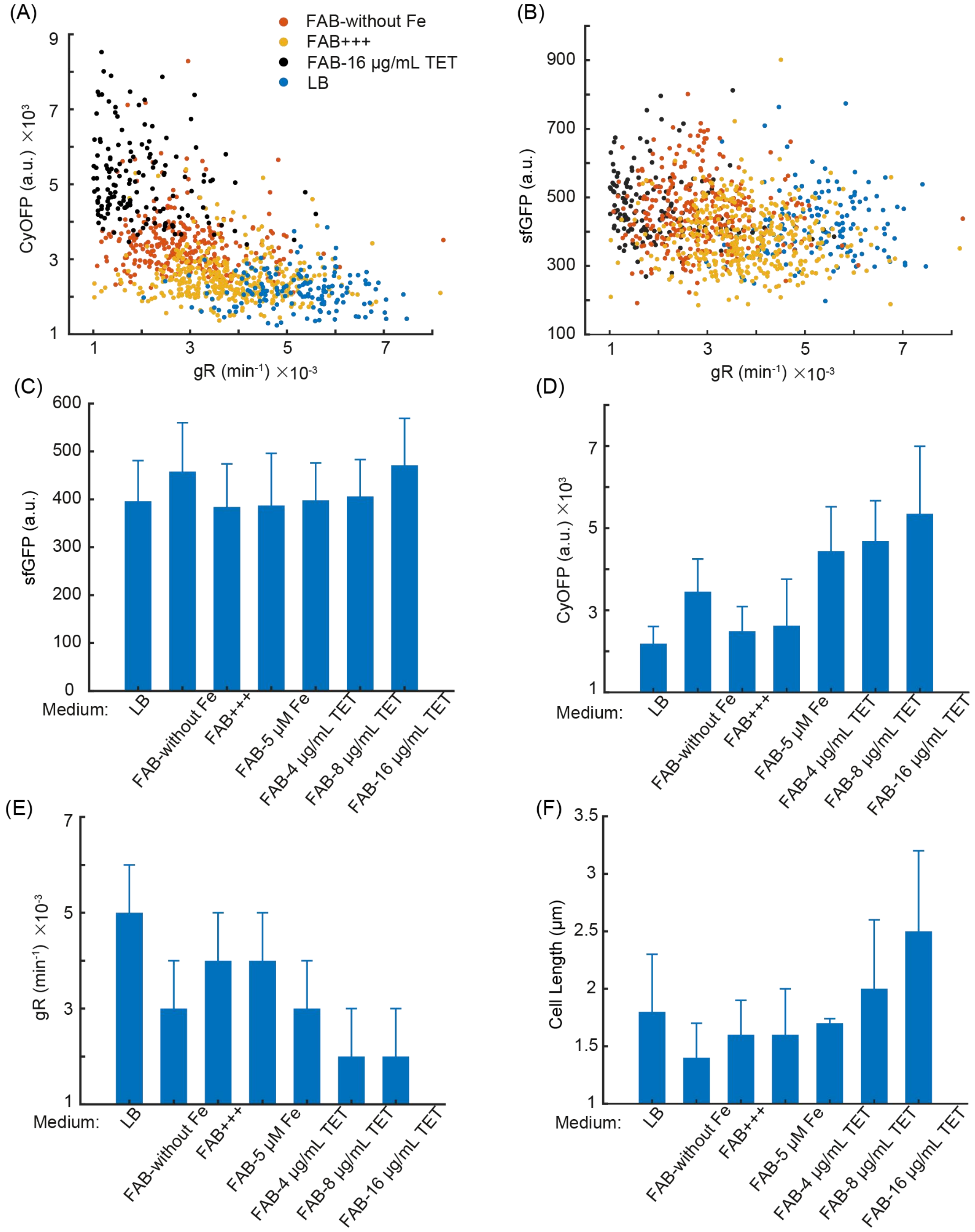


**Figure S4. Characterizing the promoter activity of the selected promoter (B173) from the promoter library at single cell level.** The measurement of the relationship between bacterial growth rate gR and the fluorescence intensity of sfGFP (A), as well as CyOFP (B), under different culture conditions. The mean values of sfGFP (C), CyOFP1 (D), bacterial growth rate (E), and bacterial length (F) were calculated under different culture conditions.

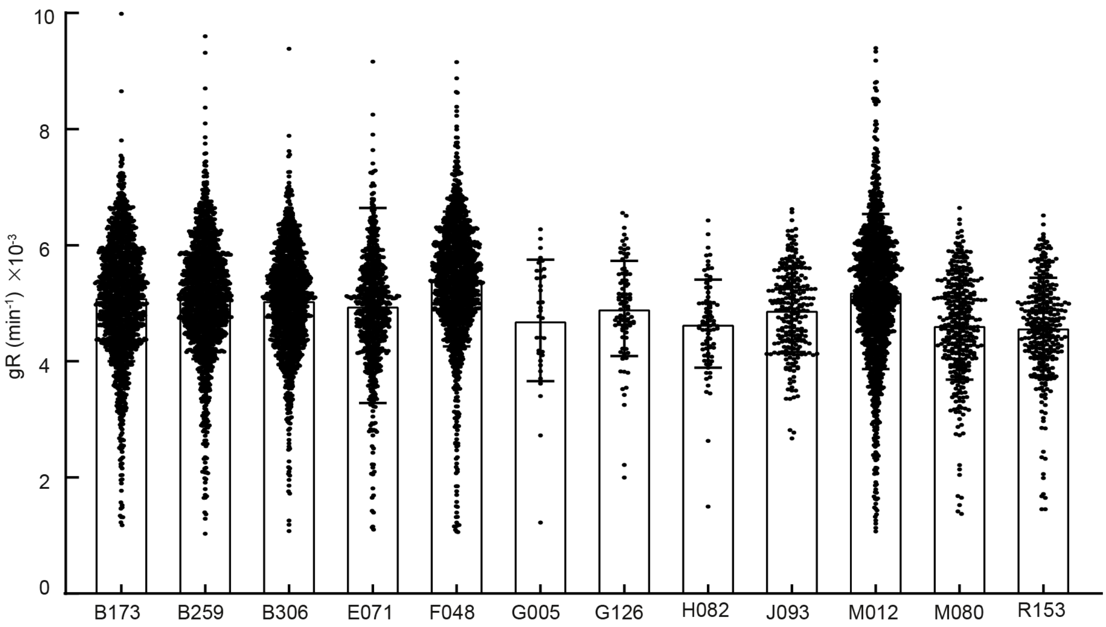


**Figure S5. Characterization of the growth rate of 12 *Pseudomonas aeruginosa* strains that contained different promoter reporting systems in LB (Luria-Bertani) medium.** The sampling interval is set to 17 min.

**TableS1. The name of bacterial strain corresponding to the promoter of gene in *Pseudomonas. aeruginosa.***

| Strains Name | Gene Name | Strains Name | Gene Name | Strains Name | Gene Name | Strains Name | Gene Name |
|---|---|---|---|---|---|---|---|
| B079 | PA1343 | E071 | PA5316 | G164 | PA0904 | N183 | PA3237 |
| B126 | PA0908 | E114 | PA4859 | G168 | PA3550 | P004 | PA3273 |
| B159 | PA2581 | E124 | PA3390 | H082 | PA2966 | P029 | PA3335 |
| B173 | PA5570 | E126 | PA3408 | H139 | PA4274 | P050 | PA3428 |
| B202 | PA1588 | E150 | PA3901 | H141 | PA4280.2 | P068 | PA3474 |
| B216 | PA3112 | E182 | PA4272 | H158 | PA4621 | P109 | PA3660 |
| B236 | PA1555 | E183 | PA4277 | H164 | PA4672 | P195 | PA3932 |
| B259 | PA4640 | F015 | PA0896 | H192 | PA5244 | Q037 | PA4100 |
| B306 | PA2975 | F048 | PA1800 | J082 | PA0386 | Q068 | PA4181 |
| B328 | PA4872 | F119 | PA3077 | J093 | PA0913 | Q072 | PA4200 |
| B352 | PA0251 | F120 | PA3119 | J139 | PA1019 | Q079 | PA4270.1 |
| B353 | PA0252 | F141 | PA3658 | J193 | PA0924 | Q084 | PA4280.5 |
| B363 | PA0267 | F210 | PA2385 | K081 | PA1147 | Q085 | PA4287 |
| B365 | PA0273 | G003 | PA4862 | K198 | PA1149 | Q137 | PA4517 |
| B423 | PA0058 | G005 | PA4918 | L040 | PA1781.1 | Q148 | PA4574 |
| B460 | PA0162 | G018 | PA5090 | L045 | PA1797 | Q154 | PA4586 |
| D071 | PA0714 | G042 | PA1786 | L073 | PA1867 | Q164 | PA4616 |
| D122 | PA0532 | G076 | PA5434 | M012 | PA2269 | Q181 | PA4684 |
| D196 | PA0680 | G094 | PA3766 | M080 | PA2484 | Q196 | PA4731 |
| D197 | PA0690 | G126 | PA3116 | N024 | PA2819 | R048 | PA4802 |
| E029 | PA3407 | G141 | PA3163 | N086 | PA2936 | R097 | PA4985 |
| E036 | PA4217 | G143 | PA4514 | N108 | PA3016 | R136 | PA5141 |
| E038 | PA3406 | G153 | PA2958.1 | N133 | PA3082 | R152 | PA5193 |
| E045 | PA5021 | G158 | PA2321 | N155 | PA3169 | R153 | PA5194 |

#### Table S2. The sequences of 12 candidate robustness promoters.

| Strains | Locus Tag | Sequence |
|---|---|---|
| B173 | *PA5570* | AACTGGTTGGACTTGCCCTCGACGAAGTTCTCGAAGGTGAAGGTACGGTTGAGATAGCTGGTGTGCTTCAGCGCACCTTCGACCTGGACGTTGCGCTCGGTGCGCACCGCAGGCGCTGCTCCAGCC<br>GGCATGGCGGCGGCCAGCGGATCGATGCTGGGTTCTTCCGGCTCGTAGGGATCGCTCGACACGCTGGGAGCCGTCGTCACTGGCGGCAGCTCTTCACGTGGCACTACCACGGGCGCGGCCGATACC<br>GGCTGCACTGGCGCCGGCGGCGGAGCAACCGGAGGCGGGGGAGCCACGTGGGTCTGCGATGGGACGATGGCGGCGCGCGGCGTACGGCTACGCTTGCTGCCTATTAATAAGGAAAGCGCGGGCAAC<br>TGACCCTCGCCGCGTTCACCGAGCAGTTCCAGAAGCCGACCGAGGTATTTCTCGTTCACCCAATCGAGGACGAAACGGTTGGGTGCATACACACGCAATTCGTCGCCTTCGGCTTCGACCTGCAAG<br>GGACGGATCCAGGTGTTGAATTGTTGGGACGGCAGCTCATCGCGGAGAAGATCCACGCACTGCTGCCAAAGTTCCACGGACACGGATATCCCCCAAAGATTGGAAAGGCGGCACAACGAAATAAGC<br>GCCTATTGTAGCCGGACGGGCCGGACTTATCCACATGCCGATGCGCTGAGGCCCCAAGCAAATCAAGACCTTAATAAAAGCAGGCAGGCCGACGGACGGTCGTCCTGGACTTTGTGGATAACCCTT<br>GCTTTGAAGGGGGTACATCCCTGCGCACAGCCGTGTGGAAAAAACCGCTGTGGATAACCATCTCGTTCATCCCCAGGCTTCGCTCAGGAAGGAAACAGCCTCGACACCGGTTCTCGACAACCTTTT<br>GCCTGCCGGAGAAACACGCAATTGCTGGGCTTCATCGAGTTATGCACAGAAAAGCGCCGTCCCAAGAGCATTAACCGTTTCTATCCTTTAAAAGAAAAATGACTTCCTTGTATCTATCTATTACGT<br>CATCGCTGGTTGGAAATTGACCTGCCCGTTCGATTTCACTAGAATCGCCGGTCTCTTTAAACGGGGTCACTGCGACCTCAAGTCGTCAACCCAGGTACCGAATCATGAAACGTACTTTCCAACCCA<br>GCACCCTCAAGCGCGCTCGCGTCCACGGTTTCCGCGCCCGTATGGCCACCAAGAACGGCCGTCAGGTTCTGTCGCGTCGTCG |
| B259 | *PA4640* | TTGCGGGTCCAGTTGTTGCGGGCTCAGGGCGCAGCCGCCGAGAACCAGGGCGCTTGCGGTCATCAGGCTGAGCAACAGGCGATGCAGCATGCTCATCTCTCCATGGGTTGGGCACGAATGGCCGGC<br>AGTATAACGGCGCCGCGAAAGCACCGCAGCATTGCCTTGGACGTTCGCCACGGGGCGCGGTTCCTCGCCTGTGGCAGGTGGCGCAGGGCACGCATGCCCGGCGTTTGGCAGATACGGCGCGGCGGA<br>TTGTCCCAGGGGGCTTTCGCCGGCGGGCACTTCGTGACCGGGAATGGGAAAAAAGTCTAAGGAGCGGGGTATTTTTTGATTTTCACCGTTATAATCGCCCCCCGATTATAAGGACGCCTCCTGATC<br>GGGTTCCGGCCCCCGCGCCATCCCCCCACAGAGAGCCGCCAGTAGAGCGCTGCCCCATGAGGCAGACGCCCTGAAACGCCTGTAGCGAGTTTTTCGGCAAGGAATGCCGCATGACCCCGCACGATT<br>CTGCGAAGGCATGCCCGCGACAGACGAGCCCTTCCGGCCCATGTCTCCAAAAGAGCGTGGATCGAAGGTATTTCACTACTTCACGAGAGTGTGGCGAGCAATGAAGAGTTTTGCGTCGGTAAGTGC<br>GCCATTAGCGCTTGAAACAGCCCCCCGCAGACGCGACCGCAGCGGATACCTCATGACCAGAGCCTGACGGCCCGTTCCGTCGCGAATGTGCTTGCACGACCGGATGCCCGCATGGGCCAGCCAGGC<br>TGCCAGATCGCCAGCCCCGCGCCCGGCATCCATAGGCGATGACGCCCGGCGTCCCGCCGTTGTTGCGAAGAATCGGACATGACGGCTCCCTCCGCCGCTGCCTGCCGGCCCGTACCGGCCTCGGCC<br>CCGCCCCGTCTGTCGTCAATTTGGTGCTGAAGATTTTGGAGAAGCGTTAATGGCGCAAAACGATCACGAAACAGTCGACATGCTGCTGGTA |
| B306 | *PA2975* | TTGTTCGGCATCAGCACCAGGTAACGGCCGGCCAGGCTGATGAAGGTGGTCAGGGCGGCGCCCTTGTTGCCGCGCTCTTCCTTCTCGACCTGGACGATGACTTCCTGGCCTTCGCTCAGGACTTCC<br>TTGATGTTGATCCGGCCTTCGGGGGATTTCTTGAAGTATTCGCGGGAGATTTCTTTGAGGGGGAGGAAGCCGTGGCGTTCGGCGCCGAAGTCGACGAAGGCGGCTTCGAGGCTGGGTTCGACGCGA<br>GTGATGCGGCCTTTGTAGATGTTGGCCTTCTTCTGCTCGCGGGCGCCCGACTCGATGTCGAGGTCGAACAGGCGTTGGCCGTCGACCAGTGCAACACGCAACTCCTCGGGTTGAGTCGCGTTGATC<br>AGCATTCTTTTCATGTAGTACCAATGGTTTCCGGGGCTACCGGAAACGGCGATCGGCACACACGACTCTCACGGTCGGTGCTCGGGCGCGTATTCAGGAATGGCGGGCCATTCCCGTGTCCAGCGG<br>CGCTCGGCACTGGGTGCCGGTCCGCGACTACGTCTCCTGCTTCCTGTGTTTCAGAAGCACTTGAGTCAGGAGGAGGAATCGGCTGTCGGAGGCGGACGAAAAATGCGTCTGGATCAAGCTAGAAGC<br>TACGCAGTCCAACAGCCGTACATCTCCACCCTACACATGTTCCTTAAGAATCGGGCGCCGCTCGCAGTATCCGGAGCGGGTTGGCATCTCGCGCTCACCGAAGTGGTCGCGGCTCATGTTTCAAGG<br>CTTTGGTTTCGAGTCGTTAGCGGGTTGGAAATTTTCGTATTTCCGCTGAAACGCCTCTGTAGGACGGCCTGACGTCCGTTCAAAGTGTTTGCAGGCGAGGGGGAAATGCAGGTTGGCAATCATCCG<br>CAGGCCAGGCCGCTTATGGCGGCTTCCCGACTATAGCAGCAATGATTAAGTGCTTCAATTGAATGAAAAATTGTTATGATGTACGGATGACTACTCCCGCCTCTCCGACCACCGGTGTCCAGCTGC<br>TCGATGTGCCGCCGGAACTGGCTGGTCAACGTATCGACAATTTCCTTATCGCCCGTCTCAAGGGTGCGCCCAAGACCCTGATCTATCGCATC |
| E071 | *PA5316* | GGAGAAAGCGGCGAGGCGCATGACGGGCTCCATTCGAACGGTTAACATGCCCATCCGCAGCGGTTGGGCGCAGTCATGCAAGGCCGTTGGCGACCTTGAATCGCTACCCTAGACCATGCGGCGGGC<br>AGCGACAACGAGGCAAGGCGACGAACGGCCGCGGTGCGAAGCCCGCAGCGAGGCGGGCCGCGGCTGCGCCATGGGCCTTCGCCCTTGCGGCGCGGGGCGCCTTTCTGATATAAAGCTGCGCTCTTT<br>TCTGGGGCCACGCTCTCCTTCGTCGGGATGCAGCGGGTAAGACCCCGGTGAAACTGAGTATCGAACTTGAGTTAAAGCAGCCAACCCATGCCGGGTTGGGCATGAGGTTATAGAGGGCTGAGGCAT<br>GTCTAGAGTCTGTCAAGTAACCGGTAAGGGTCCGGTTACCGGGAATAACATTTCCCACGCACACAACAAAACCCGCCGCCGTTTCCTGCCGAACCTGCAGCACCACCGTTTCTGGGTCG |
| F048 | *PA1800* | TGAACAGGTAGCCCTTGCCCCAGACAGTCTTGATCCGGACCGGCTCGCGCGGATTGTCCTTCAGTTTATTGCGCAGCTTGCTGATGCAGACGTCGACGCTGCGGTTCAGGCCGTCGAAACCGATGC<br>CGCGGATCTGGTTGAGGATCTCGTCGCGGGAAAGTATTTCCCCTGCCTGGCTGGCCAGCAGCCAGAGCAGCTCGAACTCCATCGTGGTCAGTTCGATGCCGAGGCCTTCGAGTTCCGCTTCGCGCC<br>GCCGTCGGTCGATGTTCAGCTTGCCGAATGTCAGGCTTTCCGGGGACGCGGGAAGGGGCGCGTGCCGGCGCATCAGGGCGCGCAGGCGGGCGAGCAGCACCGGTGGCTCGATCGGCTTGATCACGT<br>AGTCGTCGGCGCCGGACTCCAGGCCGAGGATGTGATCGAGATCGTCCTCCTGGGCGGTGAGTATGAGGATCGGCAGGTTCGCGACCCGGCGGATCTCCCGGCACACCTGCAGACCATTCTGTCCGG<br>GGAGCATGAGGTCGAGAACCACCACTTGCGGCTTGAAGTCGAGGAAGGCGGCGAACGCGGCATCACCGCGATGCACCTGGCGCACTTCGAAACCATGCTGGGAAAGGAAACTGGCGATCAGGCGGG<br>CGAGCTTCTGGTCGTCTTCGACGAGCAATACCTTGCTGAGGGTAGGGCAGTCCATAGGGTTTCATCGGTCGGAGCGGAATCGGCTGGCATTATATGCGCCCGCTTCCGGCGAAGCTCCAGCGACTC<br>GTTCGAGATCGGGCTGCGATCGGCGGAAGCGGCCGGAAACACCCCGGGGATCCGCCTGCACCCTGGCGGGGTGACTTCTGCTTTTCCCTCTACTAGAATGCATGCCCTTGATTCTGGGAGCCGGAA<br>GGACCGGCTTACGTCTGTGCAACGAGGAATATCCATGCAAGTTTCTGTTGAAAGCACTTCCGCTCTTGAGCGCCGCATGACCGTTGGTGTGCCTGCCGAGCGTATCGAGACCGAAGTCAACAAGCG<br>TCTGCAGCAGACCGCCCGTCGTGCCAAGATTCCAGGT |
| G005 | *PA4918* | ATCGGGCATCAATCCTGCTTCGATCGGCTCGCAATACACCCGCCGCGGATCGATTCGCCGGGCCAGCCGCGCCAGCCAGCGGCAGCGCAGCTCGAAATCGCCCATGCGCTTGCCGAAGGCGTGGCG<br>GTGGCTCGGCACCAGGGCGACCCGTTCGGCGCAAAGCAGCGCCCGGCGGATCACCGATTCATGGCCGGGGTGCGGCGGGTCGAACGCCCCACCATAGACCGCGAGAGCGAAGTCGGGGGCGCGTCC<br>GCCGCTCATCCCGGCAGGCAGTCGGCGAATCCGGGAACTTATTTCATCACGCATAGAAAATACCTTCCACTGCGAAATTTTGTAGTACAAGTGCTTGACGATTTATTTCACGTGATGAATTATGTA<br>CCCACATTCAGCCCCAACCCGTCAAGCGGAGGCTTCCATGAATCGCCCCAGCAAGATCGCCAGCTTCGATGTCGACGCCCAGAAAGGCTTCACCCCGCTGTGCCCGGACGAACTGCCGGTGCCCGG<br>CGGCGA |
| G126 | *PA3116* | TCAACCGCAAGGTCATCGACCAGGCTCTGGACGCCGGCACCCGCAACTACATCGGCGGCAACTGCACCGTCAGCCTGATGCTGATGGCCCTGGGCGGGCTGTTCGACGCCGGTCTGGTCGAGTGGA<br>TGAGTGCCATGACCTACCAGGCCGCCTCCGGCGCCGGCGCGCAGAACATGCGTGAACTGCTCAAGCAGATGGGCGCCGCCCATGCATCGGTGGCGGATGACCTGGCCAACCCGGCCAGCGCGATCC<br>TCGACATCGACCGCAAGGTGGCGGAGACCCTGCGCAGCGAAGCCTTCCCCACCGAGCATTTCGGCGCGCCGCTGGGCGGCAGCCTGATCCCCTGGATCGACAAGGAACTGCCCAACGGGCAGAGCC<br>GCGAAGAGTGGAAGGCCCAGGCGGAGACCAACAAGATCCTTGCCCGCTTCAAGAACCCGATCCCGGTCGACGGTATCTGCGTGCGCGTCGGCGCCATGCGTTGCCACAGCCAGGCGCTGACCATCA<br>AGTTGAACAAGGATGTGCCGCTGACCGACATCGAGGGCCTGATCAGCCAGCACAATCCCTGGGTCAAGCTGGTGCCGAACCACCGCGAGGTAAGCGTGCGCGAGCTGACCCCGGCCGCCGTCACCG<br>GCACCCTGAGCGTTCCGGTCGGGCGCCTGCGCAAGTTGAACATGGGCTCCCAGTATCTCGGCGCCTTCACCGTCGGCGACCAGTTGCTGTGGGGCGCTGCCGAGCCGCTGCGGCGCATGTTGCGCA<br>TCCTGCTGGAGCGCTGATCGATCCGGCGCCGTCCGCAATGGGCGGCGCCTCGATTGCCTGGGCGAAGGTGGGCAAGTAAAGTGCCACGCCTATTTCGCAGACCGCCGAGGCTTTTCCCATGTCCCA<br>GCCCCTCAATGTCGCAGTCGTAGGCGCCACCGGATCGGTTGGCGAGGCTCTGGTCGGTCTCCTCGACGAACGCGACTTCCCGCTTCATCGT |
| H082 | *PA2966* | AGGTCGCATTGGTAACCGGCGCCAGCCGTGGCATCGGCCAGGCGATTGCGCTGGAACTGGGGCGCCTGGGTGCCGTGGTCATCGGCACCGCGACCAGCGCGTCCGGCGCCGAGAAGATCGCCGAAA<br>CCCTCAAGGCCAATGGCGTCGAGGGTGCGGGCCTGGTCCTGGACGTTTCCAGCGACGAATCCGTAGCCGCGACCCTGGAGCACATCCAGCAGCATCTCGGCCAACCGCTGATCGTGGTCAATAACG<br>CCGGCATCACCCGCGATAATCTGCTGGTGCGCATGAAAGACGACGAGTGGTTCGATGTGGTCAACACCAACCTGAACAGTCTCTACCGTCTGTCGAAAGCCGTTCTGCGCGGTATGACCAAGGCCC<br>GCTGGGGGCGCATCATCAACATCGGTTCCGTGGTCGGCGCCATGGGCAATGCCGGGCAAACCAACTATGCCGCGGCGAAGGCCGGCCTGGAGGGCTTCACCCGTGCGCTGGCCCGGGAAGTCGGTT<br>CGCGTGCCATTACCGTGAATGCGGTGGCGCCGGGCTTCATCGACACCGACATGACCCGCGAGCTGCCGGAAGCCCAGCGCGAAGCGCTGCTGGGCCAGATTCCGCTGGGTCGCCTGGGGCAGGCGG<br>AAGAGATCGCCAAGGTGGTCGGCTTCCTCGCTTCGGACGGCGCAGCCTATGTGACCGGGGCCACCGTGCCGGTCAATGGTGGGATGTACATGAGCTGAATGTGACGTTACCTTTCTCGGGTCTGTC<br>ATAAGCGCTGTCTAGAATTCGTTGCGGAATTGCAGCCGGGTTATTAGATAGAATTATCGAAGTGGCCGTGGGAAGGTGAGGCGTTCAGCTTGAAAAGCAGAAAACCCTTTCTATACACTTACCCGC<br>AGCCCAGCTGCTCGGATTTTCCATAGGAGTGAAAACAAGGTATGAGCACCATCGAAGAACGCGTTAAGAAGATCGTCGCTGAACAACTCGGCGTGAAGGAA |
| J093 | *PA0913* | TAACGCACCTCCGCATTGTTCCCGCTGACCCGGTCCGGCTTGCCCAGCGCCCGCTCCACATCGGAGCGGGTCATTCCCGCCATTATCCGCTGCTCCACGATCGCCTTGCGGCGTTCCTTCTCGTCC<br>AGGCGTCTGCCGCAGCGATCTTCTCTTTCTCCGATCACCGCGAGCGGCACGCTTTTCCGGCCGATGGACGCCTTTTTCGTCTTCGTCTCGGCCATCGGCGTGACGCTCCTGCCTCCCGGCGGCGGG<br>TTGTCCGCCACGACGGTCTCGCCGGCCTGCCCGGCGGGGCAACCGAGTTGGGTGAAGCTGACATGGCCGCTGGCGTCTTCGCAGCGGAATACCGCAGCGGCCTCGCAGGTGGACGCCAGCGAGAGG<br>AGGAGAAGGAGGGACAGGGAACGCATGGGCGGCACTCCGGGTCGGATAAGGGTGAGTGCAGGCTAACGCGTTGCCTTTTTCCGCGGGGCCGAGTGTCCTGAGTGAAATTTCCGGCGTCGCAGTCAG<br>GCGTCGAGTGACGATGAACTTTGTGCCGCAAGCGCTTGTTCTTGCCGCCATTTTGTAACGCGCAAAACGCAAGCGGAGTGTTATCATTATCCGTATCTGCCCCGAGAGGGCCCTTGGAACACCCCC<br>ATGGACTTACCCAGTAGAAACTCGAACAGTTTCCCGTTTGACCAATCACGTTTCTGATTGACCCCTTTGGCGTGCTCCGCTACTGGGGGTGGAGCGCGCTATGACCGAAGTAGAAGCCAAGAAGCC<br>GCAGGAAA |
| M012 | *PA2269* | GCGCAGTTGGCTGGCCTCGCTGCTGGGGCTGCCGAGGATGGCCGGGGCATCGCGCAGGACGATGCGCTGGATTTCCGCTTCCAGGGCCATTTCCAGGTAGGCCCGGTTGTAGGCACGGAAGCCGCT<br>CCAGAGGTCCTCGCCGGCAGCCAGGATGGCGCCGAGGCGAAGATTCATCTCGTCGTCGATCTGCTGCACCACCGCGGCCAGCAGGCCTTTCTTGTCGCCGAAGTGGTGATACAGCGCGCCGCGGGT<br>GAGGCCGGCGCTGGCGGTCAGTTCGTCCATCGAGGCGCCGGCGTAGCCGTTGTCGGCGAACGCTCGGCGCGCGGCGTCGATCAGCTTGGCGCGGGTTTCCTCGATCATCTCGGCGCGAGGCTTGCG<br>GGCCATGCGAGGTCTCCGGGTGGCGAAGGGGCTTCAGGATAATTGACATACGCTGCGTATGTCTCCATGATTTTTGACATACGCAGCGTATGTTAATTTCGTCTTCTCCCGTCGTCGCCGGTTCCG<br>CTGGAGCCGGCCAGCCGAGGCTTTCCATGAGCAACCCCTATCTCGAACTGTTCGCCGCGCCCG |
| M080 | *PA2484* | AGGCTCACATAGGTGACGCCATAGGTCGGGGCCTTGCGGGAGCGGGCTGAGAGCATGGATGCCTCCTTGGGCTCATTTCTTCTCGCTGGTCGGCGCGGGCGCGCTGGCCTGAACCGGGCGAGTAGG<br>GGTCACTGCATTGCTGTAGAACGCGGAAGCGGCGAAGTAGCTGGCGAGGGCGCTGAAGCTGGACATGATCGGTTCTCCTGGGGAAGGGTGGGCGTCTGCCCTTGGAGAGCATGTTTGCCCAGCGGG<br>AGAACCGGCAGAAGTAAAAAGCCTGCATGGTGTATATCGAGCGCGGTGATGGATCAAGCGACCCCCTCCAGCCTTGCGACTGTTCGCCGGTGTTACCATATGACGTGTTTTTTTCTCAGGAATTCC<br>CATGGCCAAGCGTGGACGCCCCTGCGGCTTCGATCGTGAACAGGCATTGCGCCG |
| R153 | *PA5194* | ATGGAAGACCAGCGGTTGCGGCTCGAACAGGCGCACGGTTTCCTCGTGGTAGAGGCGGTGCAGGACGGTCTCGTTGTCCAGCGCCAGCAGCTCTTCGGCGGTCAGCGTGTCGGCCAGGGTGGTCAG<br>GTGCTGCCAACTGGCCTCGCGGGCCTCGGGGTCCTTCAGGCGGTCGGCGGGAAGCTGCTGCAGCAGCAGGCCGCGGGCACGGCGGCCGTTGGCGTTGAGCCAGAAACGGGTCGGCAGCTGTTCCGA<br>GCTGGCGAAGTAGTTGGACAGGCATTCCGCCAGGTTCACCCCCTCCAGGGCGACGATCCCCTGGTAGCGCTGGCCCTTGACCGGGTCCACGGTGAGGGTCAGCACGCCTTCGGGCATCAGTTCCTG<br>CATGCCGGCGTCGGCGCCGATGGACTCCGCCGAATAGCGGGCCAGGCCACGCACCTGGCGGTCGCTGGAACACTCCACCATCAACAGCGGAACCGCGCCGCTGGAGCGCGCCTGCAATACCAGCAG<br>GCCGTCGAACTTGAGCGTGCCGCAGAGCAGCGAGGCCGCCGCGAGCATCTCCCCGAGCAGCTGCGCGACCGGTTCCGGATAGGGATGTTTGGCCAGCACTTCGCTGTAGCTGCGCTCGAGATCGAC<br>CATCTCGCCGCGCACGTCGGTATCGTCGAAAAGGAAACGCTGGCTCTGATCTGAATGGGACATGCGGGACACAACGCTACGAATGAGGGAATGACAAGTGAGTGACAAAGGCCTATAAAAGCGCGC<br>TTCGTGACAGTCGTACGCCGACCCGGCGCCGATAGGGGGAAACGATTTTATGGATAATGCCTCTCCGTTCCAAGCGGCCTGGCAATGGCGCAAGTTCGCCATCTGT |

#### Table S3. Bacterial strains and plasmids used in this study.

| | Relevant characteristics | Source |
|---|---|---|
| **Strains** | | |
| PAO1 | Wild-type *Pseudomonas aeruginosa*. None resistance | J.D. Shrout |
| KT2440 | Wild-type *Pseudomonas putida*, Carb$^r$ | Laboratory storage |
| B173_PAO1 | PAO1, B173-sfGFP-J23102-CyOFP1-PUCP20. Gm$^r$ | [1] |
| B259_PAO1 | PAO1, B259-sfGFP-J23102-CyOFP1-PUCP20. Gm$^r$ | [1] |
| B306_PAO1 | PAO1, B306-sfGFP-J23102-CyOFP1-PUCP20. Gm$^r$ | [1] |
| E071_PAO1 | PAO1, E071-sfGFP-J23102-CyOFP1-PUCP20. Gm$^r$ | [1] |
| F048_PAO1 | PAO1, F048-sfGFP-J23102-CyOFP1-PUCP20. Gm$^r$ | [1] |
| G005_PAO1 | PAO1, G005-sfGFP-J23102-CyOFP1-PUCP20. Gm$^r$ | [1] |
| G126_PAO1 | PAO1, G126-sfGFP-J23102-CyOFP1-PUCP20. Gm$^r$ | [1] |
| H082_PAO1 | PAO1, H082-sfGFP-J23102-CyOFP1-PUCP20. Gm$^r$ | [1] |
| J093_PAO1 | PAO1, J093-sfGFP-J23102-CyOFP1-PUCP20. Gm$^r$ | [1] |
| M012_PAO1 | PAO1, M012-sfGFP-J23102-CyOFP1-PUCP20. Gm$^r$ | [1] |
| M080_PAO1 | PAO1, M080-sfGFP-J23102-CyOFP1-PUCP20. Gm$^r$ | [1] |
| R153_PAO1 | PAO1, R153-sfGFP-J23102-CyOFP1-PUCP20. Gm$^r$ | [1] |
| B173-CyOFP1_PAO1 | PAO1, B173-CyOFP1-PUCP20. Gm$^r$ | This study |
| B259-CyOFP1_PAO1 | PAO1, B259-CyOFP1-PUCP20. Gm$^r$ | This study |
| B306-CyOFP1_PAO1 | PAO1, B306-CyOFP1-PUCP20. Gm$^r$ | This study |
| M012-CyOFP1_PAO1 | PAO1, M012-CyOFP1-PUCP20. Gm$^r$ | This study |
| B173-RBS046/B0034 /B0030_PAO1 | PAO1, B173-RBS046/B0034/B0030-sfGFP -J23102-CyOFP1-PUCP20. Gm$^r$ | This study |
| B259-RBS046/B0034 /B0030_PAO1 | PAO1, B259-RBS046/B0034/B0030-sfGFP -J23102-CyOFP1-PUCP20. Gm$^r$ | This study |
| B306-RBS046/B0034 /B0030_PAO1 | PAO1, B306-RBS046/B0034/B0030-sfGFP -J23102-CyOFP1-PUCP20. Gm$^r$ | This study |
| M012-RBS046/B0034 /B0030_PAO1 | PAO1, M012-RBS046/B0034/B0030-sfGFP -J23102-CyOFP1-PUCP20. Gm$^r$ | This study |
| B173_KT2440 | KT2440, B173-sfGFP-J23102-CyOFP1-PUCP20. Gm$^r$ | This study |
| B259_KT2440 | KT2440, B259-sfGFP-J23102-CyOFP1-PUCP20. Gm$^r$ | This study |
| B306_KT2440 | KT2440, B306-sfGFP-J23102-CyOFP1-PUCP20. Gm$^r$ | This study |
| E071_KT2440 | KT2440, E071-sfGFP-J23102-CyOFP1-PUCP20. Gm$^r$ | This study |
| F048_KT2440 | KT2440, F048-sfGFP-J23102-CyOFP1-PUCP20. Gm$^r$ | This study |
| G005_KT2440 | KT2440, G005-sfGFP-J23102-CyOFP1-PUCP20. Gm$^r$ | This study |
| G126_KT2440 | KT2440, G126-sfGFP-J23102-CyOFP1-PUCP20. Gm$^r$ | This study |
| H082_KT2440 | KT2440, H082-sfGFP-J23102-CyOFP1-PUCP20. Gm$^r$ | This study |
| J093_KT2440 | KT2440, J093-sfGFP-J23102-CyOFP1-PUCP20. Gm$^r$ | This study |
| M012_KT2440 | KT2440, M012-sfGFP-J23102-CyOFP1-PUCP20. Gm$^r$ | This study |
| M080_KT2440 | KT2440, M080-sfGFP-J23102-CyOFP1-PUCP20. Gm$^r$ | This study |
| R153_KT2440 | KT2440, R153-sfGFP-J23102-CyOFP1-PUCP20. Gm$^r$ | This study |

| **Plasmids** | | |
|---|---|---|
| B173-CyOFP1-PUCP20 | Plasmid used for monitoring B173 promoter expression using CyOFP1 as a reporter gene. $Gm^r$ | This study |
| B259-CyOFP1-PUCP20 | Plasmid used for monitoring B259 promoter expression using CyOFP1 as a reporter gene. $Gm^r$ | This study |
| B306-CyOFP1-PUCP20 | Plasmid used for monitoring B306 promoter expression using CyOFP1 as a reporter gene. $Gm^r$ | This study |
| M012-CyOFP1-PUCP20 | Plasmid used for monitoring M012 promoter expression using CyOFP1 as a reporter gene. $Gm^r$ | This study |
| B173-RBS046/B0034/B0030-sfGFP-J23102-CyOFP1-PUCP20 | Plasmid was used for characterizing whether RBS affected the robustness of B173 promoter by replacing RBS upstream of sfGFP with RBS046 or B0034 or B0030. $Gm^r$ | This study |
| B259-RBS046/B0034/B0030-sfGFP-J23102-CyOFP1-PUCP20 | Plasmid was used for characterizing whether RBS affected the robustness of B259 promoter by replacing RBS upstream of sfGFP with RBS046 or B0034 or B0030. $Gm^r$ | This study |
| B306-RBS046/B0034/B0030-sfGFP-J23102-CyOFP1-PUCP20 | Plasmid was used for characterizing whether RBS affected the robustness of B306 promoter by replacing RBS upstream of sfGFP with RBS046 or B0034 or B0030. $Gm^r$ | This study |
| M012-RBS046/B0034/B0030-sfGFP-J23102-CyOFP1-PUCP20 | Plasmid was used for characterizing whether RBS affected the robustness of M012 promoter by replacing RBS upstream of sfGFP with RBS046 or B0034 or B0030. $Gm^r$ | This study |

**Table S4. These core sequences were exhibited in detail for the four selected promoters (B306, M012, B259, B173).**

| **Strains** | **Locus Tag** | **Sequence** |
|---|---|---|
| B306 | Pro-F7 | TTCCCGACTATAGCAGCAATGATTAAGTGCTTCAATTGAATGAAAAATTGTTATGATGTACGG |
| M012 | Pro-F5 | AGGGGCTTCAGGATAATTGACATACGCTGCGTATGTCTCCATGATTTTTGACATACGCAGCGTATGTTAATTTCGTCTTCTCCCGTCGTCGCCGGTTCCGCTGGAGCCGGCCAGCCGAGGCTTTCC |
| B259 | Pro-F3R4 | AAAAAGTCTAAGGAGCGGGGTATTTTTTGATTTTCACCGTTATAATCGCCCCCCGATTATAAGGACGCCTCCTGATCGGGTTCCGGCCCCCGCGCCATCCCCCCACAGAGAGCCGCCAGTA |
| B173 | Pro-F6 | CAATTGCTGGGCTTCATCGAGTTATGCACAGAAAAGCGCCGTCCCAAGAGCATTAACCGTTTCTATCCTTTAAAAGAAAAATGACTTCCTTGTATCTATCTATTACGTCATCGCTGGTTGGAAATTGACCTGCCCGTTCGATTTCACTAGAATCGCCGGTCTCTTTAAACGGGGTCACTGCGACCTCAAGTCGTCAACCCAGGTACCGAATC |

**TableS5. Arrangement of strains in high throughput microscopic experiments.**

| | | | | | | | | | | | |
|---|---|---|---|---|---|---|---|---|---|---|---|
| 1：<br>R136 | 2：<br>B236 | 3：<br>B423 | 4：<br>E036 | 5：<br>E182 | 6：<br>G003 | 7：<br>G143 | 8：<br>H158 | 9：<br>K198 | 10：<br>N108 | 11：<br>P109 | 12：<br>Q137 |
| 24：<br>R152 | 23：<br>B259 | 22：<br>B460 | 21：<br>E038 | 20：<br>E183 | 19：<br>G005 | 18：<br>G153 | 17：<br>H164 | 16：<br>L040 | 15：<br>N133 | 14：<br>P195 | 13：<br>Q148 |
| 25：<br>B079 | 26：<br>B306 | 27：<br>R153 | 28：<br>E045 | 29：<br>F015 | 30：<br>G018 | 31：<br>G158 | 32：<br>H192 | 33：<br>L045 | 34：<br>N155 | 35：<br>Q037 | 36：<br>Q154 |
| 48：<br>B126 | 47：<br>B328 | 46：<br>D071 | 45：<br>E071 | 44：<br>F048 | 43：<br>G042 | 42：<br>G164 | 41：<br>J082 | 40：<br>L073 | 39：<br>N183 | 38：<br>Q068 | 37：<br>Q164 |
| 49：<br>B159 | 50：<br>B352 | 51：<br>D122 | 52：<br>E114 | 53：<br>F119 | 54：<br>G076 | 55：<br>G168 | 56：<br>J093 | 57：<br>M012 | 58：<br>P004 | 59：<br>Q072 | 60：<br>Q181 |
| 72：<br>B173 | 71：<br>B353 | 70：<br>D196 | 69：<br>E124 | 68：<br>F120 | 67：<br>G094 | 66：<br>H082 | 65：<br>J139 | 64：<br>M080 | 63：<br>P029 | 62：<br>Q079 | 61：<br>Q196 |
| 73：<br>B202 | 74：<br>B363 | 75：<br>D197 | 76：<br>E126 | 77：<br>F141 | 78：<br>G126 | 79：<br>H139 | 80：<br>J193 | 81：<br>N024 | 82：<br>P050 | 83：<br>Q084 | 84：<br>R048 |
| 96：<br>B216 | 95：<br>B365 | 94：<br>E029 | 93：<br>E150 | 92：<br>F210 | 91：<br>G141 | 90：<br>H141 | 89：<br>K081 | 88：<br>N086 | 87：<br>P068 | 86：<br>Q085 | 85：<br>R097 |

**Table S6. The truncated B306 promoter, consisting of only 63 bases, was evaluated in chassis cells with various regulatory factors knocked out.**

| | sfGFP_B306_Pro-F7 | | CyOFP1_B306_Pro-F7 | |
|---|---|---|---|---|
| | Stationary phase | logarithmic phase | Stationary phase | logarithmic phase |
| **PAO1_wt** | 208 | 190 | 4709 | 2438 |
| **Delt-OxyR** | 203 | 182 | 3022 | 1977 |
| **Delt-RpoF** | 219 | 189 | 4724 | 2286 |
| **Delt-RpoS** | 220 | 191 | 4973 | 2507 |
| **Delt-RpoN** | 210 | 183 | 3716 | 1756 |

The expression of sfGFP, serving as a reporter gene for the promoter, was measured, while CyOFP1 expression driven by the constitutive promoter J23102 served as a control. The table below presents the average fluorescence intensity obtained from three experimental replicates.

### REFERENCES:

1. Chen W, Zhang J, Li F, Wang C, Zhang Y, et al. Genome-Wide Analysis of Gene Expression Noise Brought About by Transcriptional Regulation in Pseudomonas aeruginosa. *mSystems*. 2022, *0* (0), e00963-22.
